# Overexpression of DnaJ chaperones in ameliorating toxicities associated with FUS and TDP-43

**DOI:** 10.1101/2025.05.28.656528

**Authors:** Snehal Ahire, Tania Bose

## Abstract

Amyotrophic Lateral Sclerosis (ALS) is an adult-onset neurodegenerative disease. It primarily affects the motor neurons, leading to muscle weakness and eventually death of the patients. Various factors, such as genetics, environment, age, etc, are involved in the etiopathogenesis of ALS. As ALS is a highly challenging and complex disease involving various pathogeneses linked with progressive motor neuron degeneration, it is difficult to have a single therapeutic target against this disease. ALS mutants of TDP-43 and FUS are showing slow growth as well as protein aggregation in the cytoplasm in yeast. Our study involves cloning of 38 *Drosophila* DnaJ domain chaperones and overexpressing them in a yeast ALS model. After the interaction of mutants TDP-43 and FUS with the DnaJ domain chaperones, they were categorized into suppressors and enhancers. The major hits on our screen were CG7872, Hsc70-4, and Mrj, which rescued TDP-43 toxicity by reducing protein aggregation and slow growth in yeast cells. In FUS, slow growth and protein aggregation are reduced with the chaperones CG32641 and CG6693, resulting in a lowering of FUS toxicity. On the other hand, chaperones CG2911, CG2790, and CG8476 in presence of TDP-43 overexpression, grows slower with an increase in the number of aggregates in yeast cells. These chaperones increase the toxicity of TDP-43. Similarly, in FUS, Hsc20, CG7130, CG2911, and JdpC also showed slow growth phenotype and an increase in protein aggregation in yeast cells. These chaperones enhance the toxicity of FUS. This screening will enhance our understanding of the ALS pathology, and the chaperones identified in this study can serve as neuroprotective agents for ALS.

**Graphical abstract:** 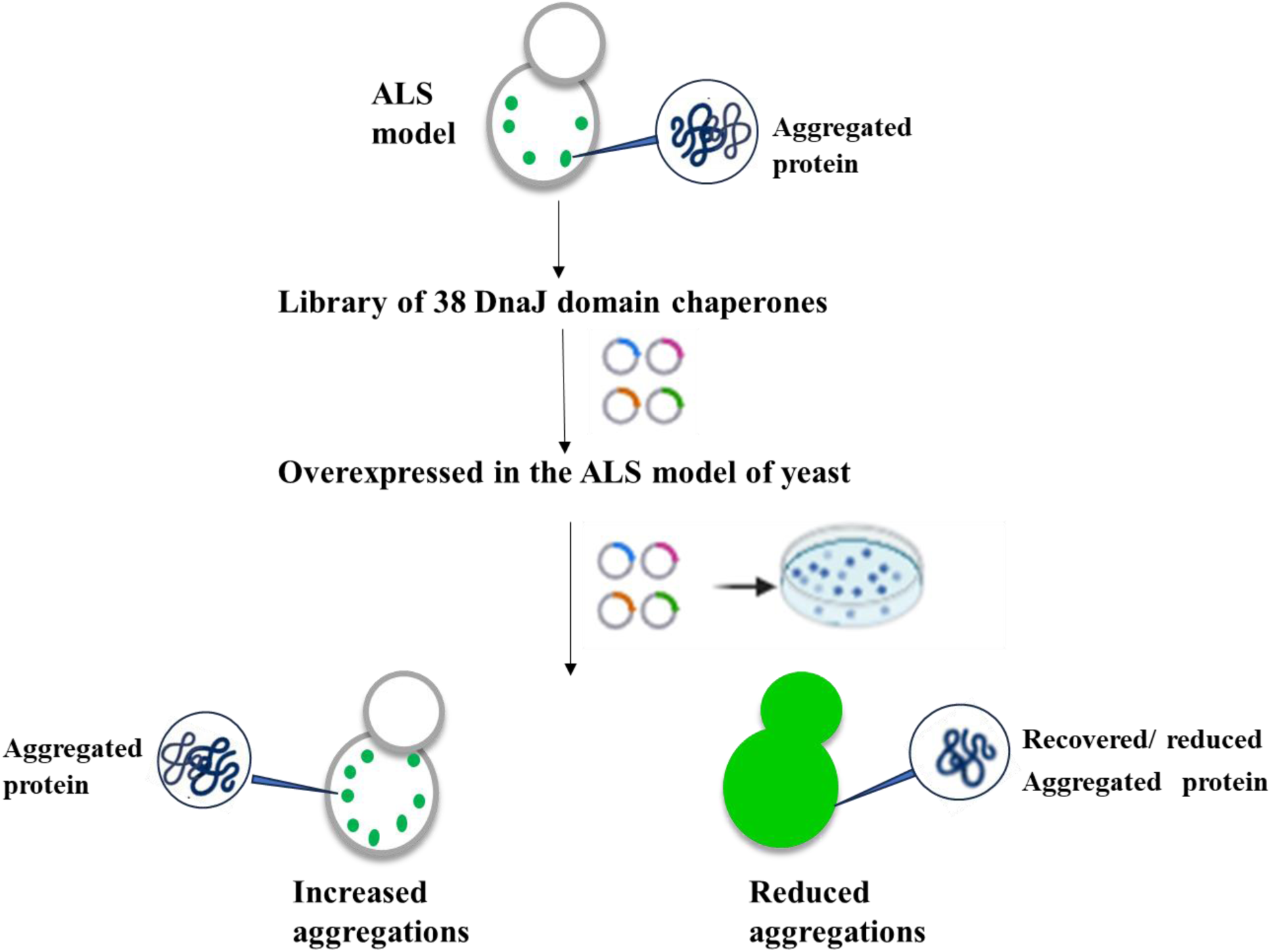

## 1.0 Introduction

ALS involves an interplay of different epigenetic genes associated with its pathogenesis. These genes include SOD1, FUS, TARDBP (TDP-43), VAPB, VCP, and OPTN (Cirulli et al., 2015; Ghasemi & Brown, 2018; J. O. Johnson et al., 2010; Rosen et al., 1993). These genes are found in all types of pathophysiology associated with ALS. ALS is found in two forms: sporadic and familial. ALS also results from family history and lifestyle factors, including smoking and dietary factors (Armon., et al 2009; Jong et al., 2012).

From our study, we focus on the cellular toxicity and protein aggregation phenotypes of ALS (B. S. Johnson et al., 2008a). In this study, two ALS mutants, TDP-43 and FUS, when transformed with 38 DnaJ domain chaperones, show either slow or enhanced growth combined with minimal or increased in protein aggregation, respectively.

Studies from other reports reveal the presence of pathogenic protein aggregates in mutant SOD-1 ALS cases (Watanabe et al., 2001). Protein aggregation is due to the failure of protein quality control system (PQC) failure to handle folding and the failure of degradation of misfolded proteins. (Mogk & Bukau, 2006). Protein aggregation is not always toxic. Sometimes, protein aggregation can be a protective storage mechanism. However, within the cell, depending on the location and size of protein aggregates, these may be considered toxic (Kampinga et al., 2016)

Neurodegenerative diseases like ALS, Alzheimer’s, Huntington’s, and Parkinson’s have a protein aggregation trait, a specific protein that can fold into an alternative stable form, leading to protein aggregation and accumulation as fibrillar deposits in tissues (Carrell & Lomas, 1997; Goedert, 2004; Martindale et al., 1998).RNA processing pathways and RNA-binding proteins are involved in the etiopathogenesis of ALS (Gitler & Shorter, 2011a; Lagier-Tourenne & Cleveland, 2009). Many of the genes associated with ALS pathogenesis are RNA-binding protein (RBP) genes, including TDP-43, FUS, Ataxin-2, TAF15, EWSR1, and matrin3 (Couthouis et al., 2011; Elden et al., 2010a; Malik et al., 2018; Ticozzi et al., 2011). All these genes share structural similarities, including RNA-binding domains, and exhibit some functional similarities, as they are involved in RNA metabolism and localize in stress granules.

TDP-43 and FUS have RNA-binding domains, and after mutation, both genes have been reported to be toxic in multiple studies. In 2006, Arai and Newmann found TDP-43 to be an essential component of ubiquitinated protein aggregates in FTLD-U and sporadic ALS (Arai et al., 2006). In ALS, when TDP-43 is derived from the damaged central nervous system, toxic TDP-43 is found in hyperphosphorylated form, cleaved to obtain a C-terminal fragment, and ubiquitinated (Neumann et al., 2006a). FUS also exhibits ubiquitous expression, similar to TDP-43, and its expression is independent. After the missense mutation, FUS also forms cytoplasmic aggregates in neurons (Vance, Rogelj, Hortobágyi, et al., 2009). In 0.4% of familial ALS cases, FUS/TLS mutations were found (Kwiatkowski et al., 2009a; Vance, Rogelj, Hortobágyi, et al., 2009). Although TDP-43 and FUS have RNA-binding domains and similar cytoplasmic aggregation properties, their toxicity mechanisms have very subtle differences (Sun et al., 2011).

We used *Drosophila* DnaJ domain chaperones, i.e Hsp40, which help in protein folding. These are the proteins that help chaperones in protein folding. These -binding molecules help to fold the proteins by binding Hsp70 or Hsp90, or both. DnaJ chaperones can induce or inhibit the ATP hydrolysis activity of Hsp70 and Hsp90 chaperones(Bar-Lavan et al., 2016; Tsai & Douglas, 1996). There are different Hsp40s in mammals, many of which follow the categories based on their domain architecture (Cheetham & Caplan, 1998). Most of them are classified into two groups: J domain proteins and tetracopeptide repeats, i.e., TPR (Caplan, 2003). Together, they form a network to modulate the functions, identifies a subset of misfolded proteins which acts as a substrate and interacts with other chaperones and co-chaperones (Caplan, 2003).

There have been reports using DnaJ library in yeast to investigate protein aggregation in ALS. We further attempted this study using an overexpression screening to accompany the earlier studies to answer the aggregation and disaggregation properties of DnaJ domain chaperones. Multiple studies have shown that heat shock proteins and chaperones help to reduce the toxicity of mutant TDP-43 as well as FUS. In HEK293 cells, overexpression of HSF-1/DNAJB21/HSP70 shows reduced cytoplasmic protein aggregation caused by mutant TDP-43 (Vance, Rogelj, Hortobágyi, et al., 2009). Previous work has shown that HSP40 chaperone Sis1, plays a role in protein folding, transport of misfolded proteins, and catabolism of ubiquitin-dependent misfolded proteins. When Sis1 is overexpressed in TDP-43 cells, it lowers the toxicity of TDP-43. Its mammalian homolog, DnaJB1, when overexpressed in rodent primary cortex, also lowers TDP-43 toxicity in neurons (Park et al., 2017). In HeLa cell line, it was observed that HSP70 prevents amyloid aggregation caused by the mutant TDP-43 and prevents the liquid-to-solid transition (Gu et al., 2021).

There are very few studies that show HSPs can increases TDP-43 toxicity, one in vivo study on *C. elegance* showing that HSP 90 chaperone contributes to increase in TDP-43 toxicity (Garcia-Toscano et al., 2024).

TDP-43 and FUS both undergo liquid-to-solid phase separation (Li et al., 2018; Molliex et al., 2015). This process results in the formation of liquid-like sections in the cytoplasm during stress and at the site of DNA damage. The pHSP 40 chaperones Hdj2 (DNAJA1) and Hdj1 (DNAJB1) assist FUS in undergoing phase separation. Additionally, Hdj1 aids the phase-separated FUS in inhibiting amyloid and protein aggregation within the cell(Gu et al., 2020). The small heat shock protein Hsp27 is co-localized with FUS and prevents their phase separation by binding FUS to the low complexity region, thereby helping to avoid the formation of stress granules in the cell model (Z. Liu et al., 2020).

Chaperones are the proteins that assist in the folding or unfolding of large proteins and differentiate between misfolded proteins, unfolded proteins, and aggregated proteins with specific biological functions (Beissinger & Buchner, 1998). The main cellular function of the chaperones is assisting in the quality control of protein folding. Not all chaperones can prevent aggregation. Some chaperones assist in restoring correct folding and denatured protein function and aid in the assembly and disassembly of proteins. They also help fold *de-novo* proteins and, in the translocation and disaggregation of protein aggregates (Bar-Lavan et al., 2016). Chaperones are classified into five main families based on their molecular mass and sequence similarities (Bascos & Landry, 2019).

In prokaryotes and eukaryotes, Heat shock proteins (HSPs) are ubiquitously conserved for the survival of stressful conditions like heat stress (Morimoto & Santoro, 1998). It can cause harmful damage to cellular architecture and interfere with vital cellular functions. To combat heat stress, the cell activates its classical pathway, whereby it expresses heat shock proteins for a shorter period for cell survival and maintains protein homeostasis (Lindquist, 1981).

In our study, we have used the yeast model to focus on the mechanism of cellular toxicity and protein aggregation phenotype in both TDP-43 and FUS (B. S. Johnson et al., 2008b, 2009; Sun et al., 2011). Our observation supports enhanced and rescued protein aggregation and cellular growth via interaction of DnaJ domain chaperones with ALS mutants. We found potential suppressors and enhancers with the interaction of DnaJ domain chaperones. To check our findings, we expressed 38 DnaJ domain chaperones from flies in yeast (Deo et al., 2024; Desai et al., 2024).

Gitler and colleagues established yeast as a model to study ALS related to TDP-43 [(B. S. Johnson et al., 2008a). The same group also reported using yeast for genome-wide screening of FUS to uncover the mechanisms of the cellular pathways and the genes involved in FUS toxicity (Sun et al., 2011). Taking cues from these studies, we explore the interactions between TDP-43 and FUS with Drosophila DnaJ domain chaperones, including their potential suppressors and enhancers.

This study focuses on two genes associated with ALS, namely TDP-43 and FUS. Despite extensive research, there is still limited information available about its cure and precise cause. In this work, we discuss screening both genes by transforming with *Drosophila* DnaJ domain chaperones and identifying potential suppressors and enhancers. This will help identify the possible interactors and their network with respect to the DnaJ domain chaperones.

Our study emphasizes protein aggregation and cellular toxicity in ALS mutants, involving the overexpression of TDP-43 and FUS genes. We screened 38 DnaJ domain chaperones of *Drosophila*, which include CG30156, CG5001, Mrj, DnaJ60, CG2911, CG6693, JdpC, CG9828, CG4164, CG2887, CG12020, JdpA, CG10565, Tpr2, Csp, CG7394, CG7130, CG8476, CG7872, Hsc20, Sec63, P58IPK, CG43322, CG7133, CG7556, CG14650, CG32641, CG11035, CG10375, CG17187, DnaJ1, CG3061, CG8531 and CG7387. In addition, there are, two Hsc70 (Hsc70-1, Hsc70-4) and Hsp70 (Hsp70Aa and Hsp70Bb) chaperones. We chose only those chaperones that play a role in protein folding, listed in **Supplementary Table S2**. Their function and activity are the same as the human homologs (Gaudet et al., 2011; R. et al., 2012).

We found that chaperones CG7872, Mrj, and Hsc70-4 suppress the toxicity of TDP-43, and chaperones CG32641 and CG6693 suppress the toxicity of FUS.

These chaperones are found to play a protective role in other disease conditions as well. DNAJC25 is an orthologue of CG7872, localised in the cytoplasm, and plays a role in hepatocarcinoma suppression (LIU et al., 2012a).

After mutation, Mrj is found in disease conditions like frontotemporal dementia and limb-girdle muscular dystrophy type 1(Yabe et al., 2014). In some cases, in mouse and *Drosophila*, Mrj inhibits Huntington PolyQ aggregation [48,36]. In addition to Huntington, Mrj also reduces the α-synuclein aggregation in Parkinson’s disease with co-chaperone Hsp70 (Aprile et al., 2017). In both Huntington’s and Parkinson’s disease, Mrj suppress the toxicity of Htt and α-synuclein disease-causing proteins, but no such evidence has been found in the ALS-causing gene TDP-43. Also, Hsc70 is a co-chaperone of DnaJ domain chaperones and is also found to improve the TDP-43 toxicity in sporadic ALS patients (Arosio et al., 2020; Richter et al., 2010). Human orthologue of Hsc70-4 is HSPA8 which has been reported to be involved in clearance of misfolded proteins (T. Liu et al., 2012; Stricher et al., 2013).

DNAJC9 is the human orthologue of CG6693 and has been found to play a role in cancer (Zhang et al., 2020). Also takes part in histone chaperone function with the help of the co-chaperone Hsp70 (Hammond et al., 2021). DNAJB9, is also a human orthologue of chaperone CG32641 and is found associated with ER degradation machinery, and also interacts with CFTR and rescues the cystic fibrosis-causing gene Δ F508 (Huang et al., 2019; Lai et al., 2012). Some chaperones can enhance the toxic phenotype, specifically slow growth and protein aggregation in both mutant FUS and TDP-43. We found that chaperones CG2911, CG2790, and CG8476 are enhancers of TDP-43 toxicity, while chaperones Hsc20, CG2911, CG7130, and JdpC enhances FUS toxicity. Previously, they were found to contribute to toxicity in certain diseased conditions (D. Liu et al., 2024; Thomson & Dinger, 2016; Uhrigshardt et al., 2013; Veenma et al., 2018).

To develop TDP-43 and FUS models in *Saccharomyces cerevisiae,* we used galactose-inducible plasmids pRS426 Gal-TDP-43-GFP and pAG416 Gal-FUS-YFP, respectively. Mutant TDP-43 and FUS shows slow growth and protein aggregation in yeast cells at permissive temperature. When these two mutants were expressed with 38 DnaJ domain chaperones, we found some chaperones suppressed the toxicity, while some chaperones enhance the toxicity of the mutants, TDP-43 and FUS.

## 2.0 Results

### 2.1 Yeast model of TDP-43 and FUS: growth assay and protein aggregation

We constructed TDP-43 and FUS yeast models with TDP-43 and FUS plasmids pRS426 Gal-TDP-43-GFP and pAG416 Gal-FUS-YFP, respectively [method adopted from (Deo et al., 2024; Desai et al., 2024). This construct has a Ura marker with a GFP tag at the C-terminal and a galactose-inducible promoter. **(Supplementary Fig S1)**

The TDP-43 mutant results in slow growth and protein aggregation in yeast cells, indicating pathogenicity (B. S. Johnson et al., 2008a). Similarly, the FUS mutant shows slow growth and protein aggregation, indicating pathogenicity in yeast cells (Sun et al., 2011).

We used an empty vector as a control, and TDP-43 and FUS for determining ALS toxicity (Mogk, 1999; Tomoyasu et al., 2001). The spot dilution assay of TDP-43 and FUS mutants shows slow growth compared to the control in Gal-Ura **(Supplementary Fig S2a).**

A quantitative study of both mutants was done using a 48-hour growth assay. TDP-43 and FUS cultures were added in triplicates to a 96-well plate at 30°C with orbital shaking. Growth assay of TDP-43 and FUS shows slowdown of TDP-43 and FUS growth in liquid medium, i.e, Gal-Ura, compared to the control **(Supplementary Fig S2b).**

We further checked whether our mutants show protein aggregation under confocal microscopy. For the microscopy, cultures were inoculated in SD-Ura, on the following day. After raffinose wash and addition of 20% Galactose, we find two morphologies in confocal microscopy: diffuse cytoplasmic and punctae forms. In TDP-43 and FUS, we find more punctae, indicating protein aggregation in the yeast cell, and in control, we observe diffuse cytoplasmic signal, which indicates no protein aggregation in the cell **(Supplementary Fig S2c).**

Since, TDP-43 and FUS mutant strains show a slow growth phenotype and aggregation of proteins compared to control, we further checked these mutants against 38 DnaJ domain chaperones to determine, how it enhances or rescues growth and aggragation properties.

### 2.2 Classification of enhancers and suppressors of TDP-43 and FUS-*Drosophila* DnaJ domain chaperones, based on growth assay

We transformed 38 DnaJ domain chaperones ALS mutants, TDP-43 and FUS. These transformed colonies of DnaJ domain chaperones were selected and inoculated in the SD-Trp-Ura medium. On the second day, cultures were spotted onto SD-Trp-Ura and SGal-Trp-Ura plates. SD-Trp-Ura is the control, and SGal-Trp-Ura serve as a test. Chaperones that enhance the effect of the mutation are collectively called enhancers. Enhancers are those whose growth are reduced in comparison to control TDP-43 and FUS. Since the plasmids are galactose inducible, we find that chaperones CG2911, CG2790 and CG8476 grow slower on SGal-Trp-Ura plates than control TDP-43; hence, these are enhacers of TDP-43 toxicity (**Fig. 1a).** Chaperones Hsc20, CG2911, JdpC grow slower in SGal-Trp-Ura plates than in FUS with empty vector strain. These chaperones are enhancers of FUS (**Fig. 1f).**

**Table 1.** List of weak enhancers and no effect chaperones on ALS mutant TDP-43 and FUS.

| TDP-43 |  | FUS |  |
| --- | --- | --- | --- |
| Weak enhancer | No effect | Weak enhancer | No effect |
| CG12020 | CG14650 | CG9828 | CG7556 |
| CG7394 | Tpr2 | CG12020 | CG17187 |
| CG4164 | CG7556 | Hsp70Aa | P58IPK |
| Dnaj60 | CG9828 | Hsp70Bb | Csp |
| CG3061 | Hsp70Bb | CG11035 | Sec63 |
| Hsp70Aa | P58IPK | CG3061 | CG43322 |
| CG32641 |  | Dnaj60 | CG11035 |
| DnaJ1 |  | CG7133 | Hsc70-1 |
| JdpC |  | CG3061 | CG10565 |
| CG2887 |  | CG14650 |  |
| Hsc20 |  | CG2887 |  |
| JDPA |  | CG10375 |  |

**Fig. 1.**
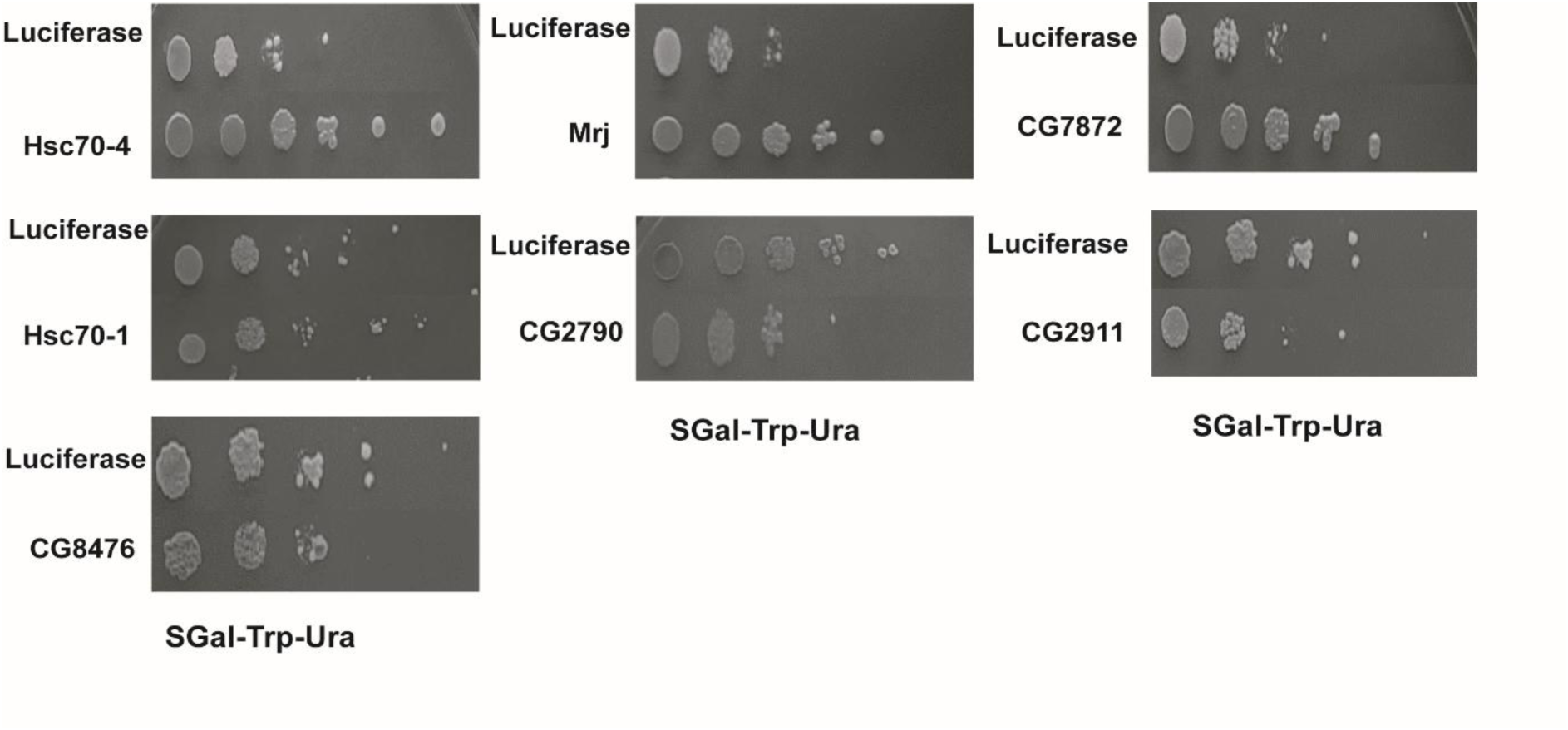
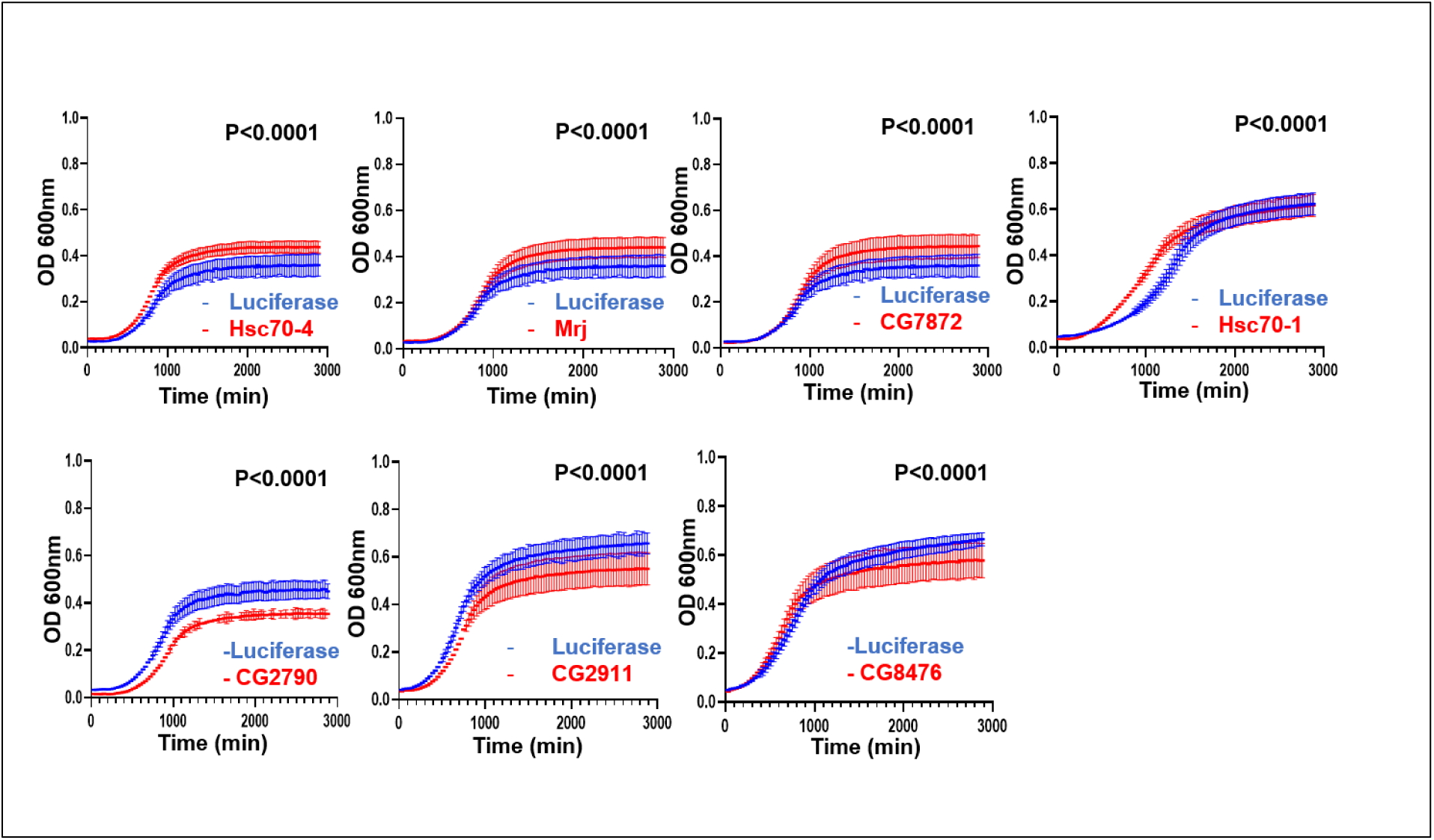
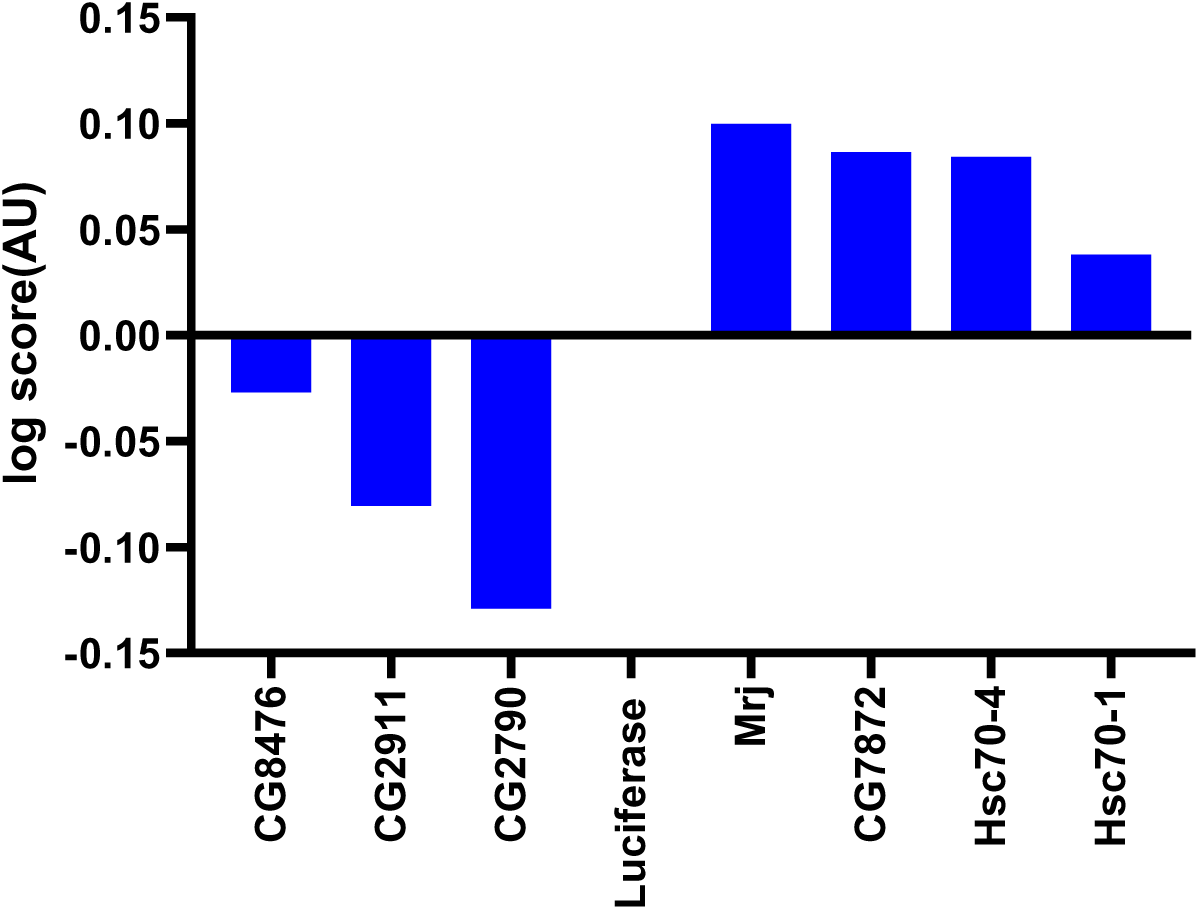
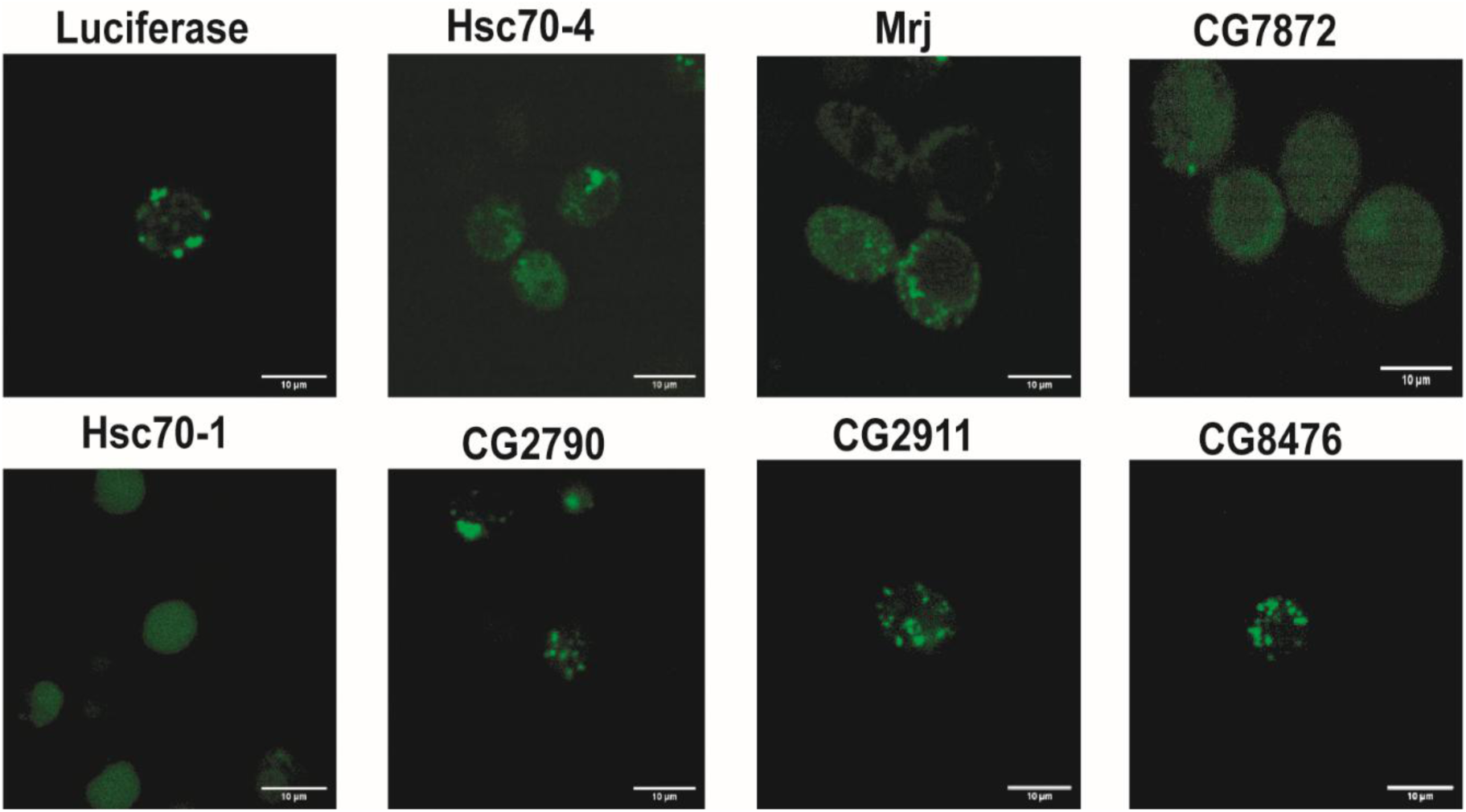
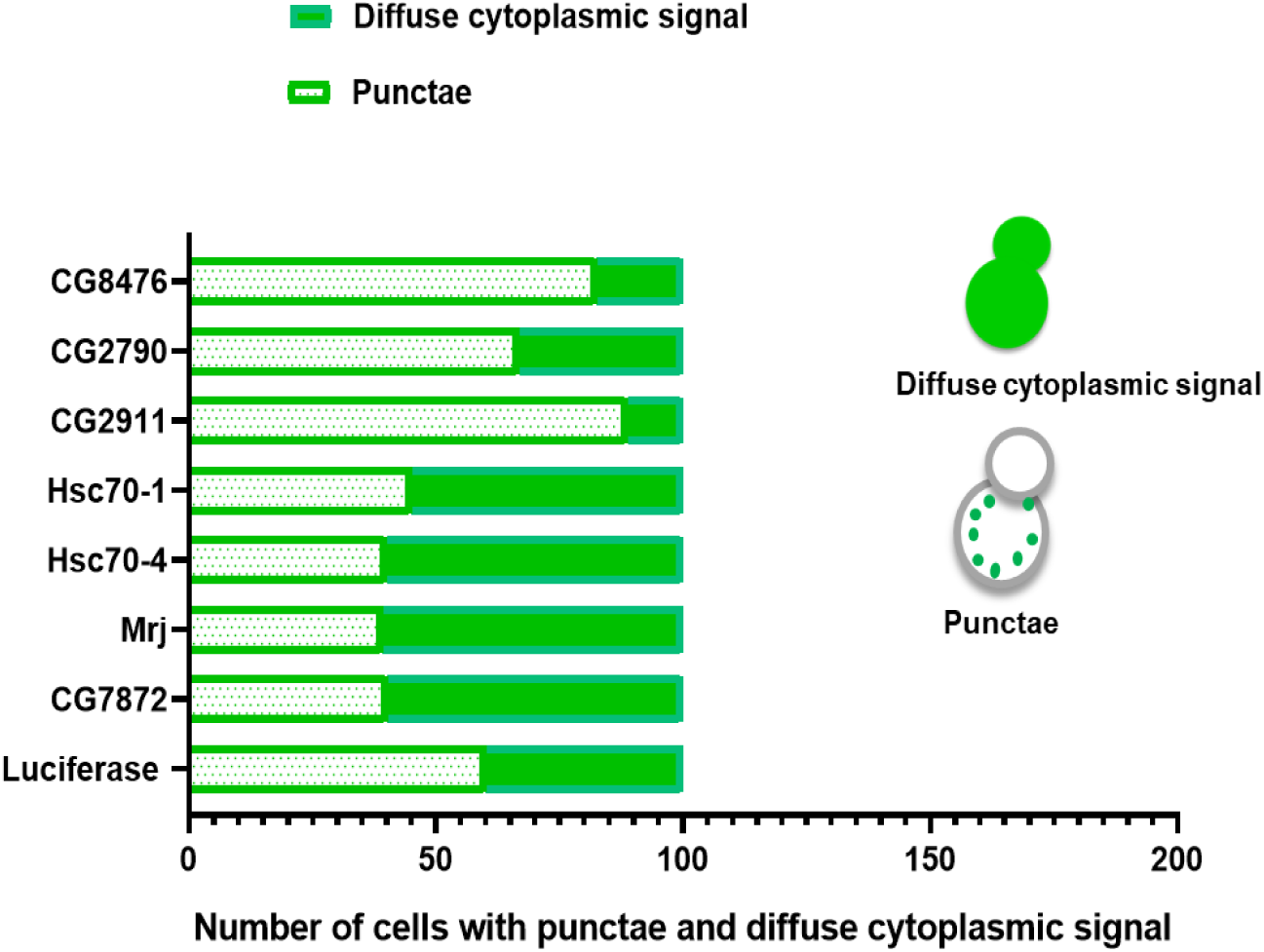
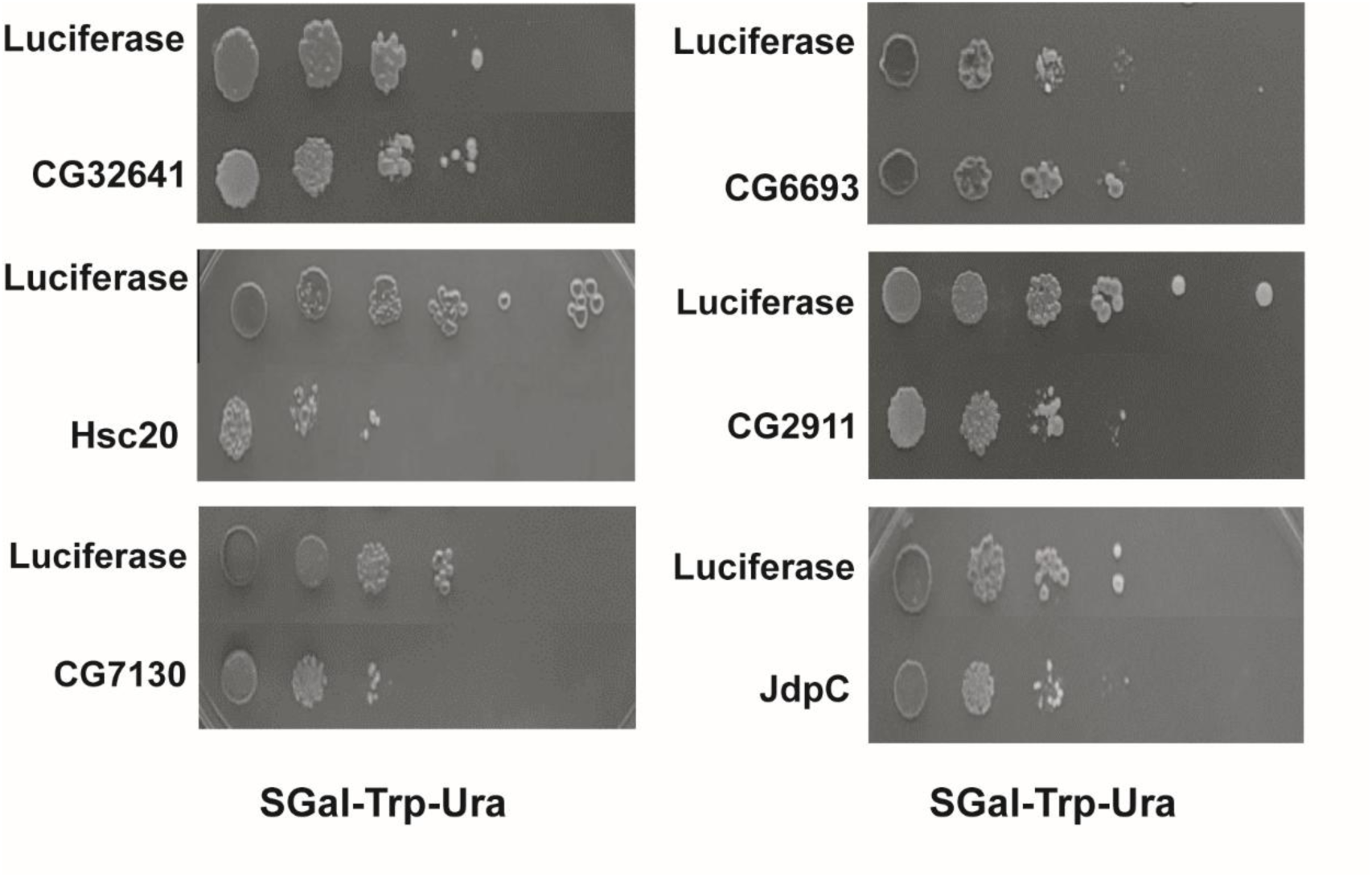
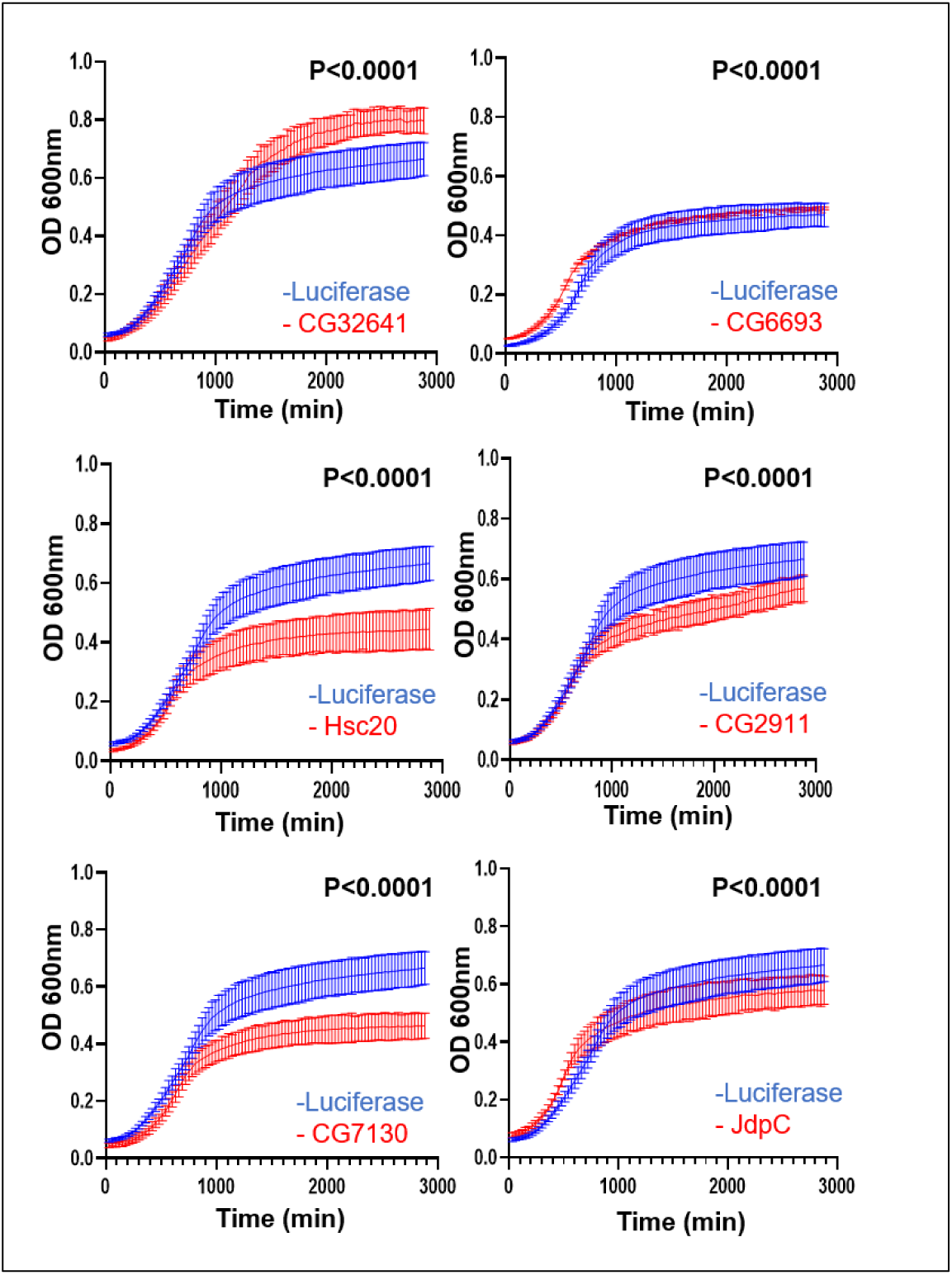
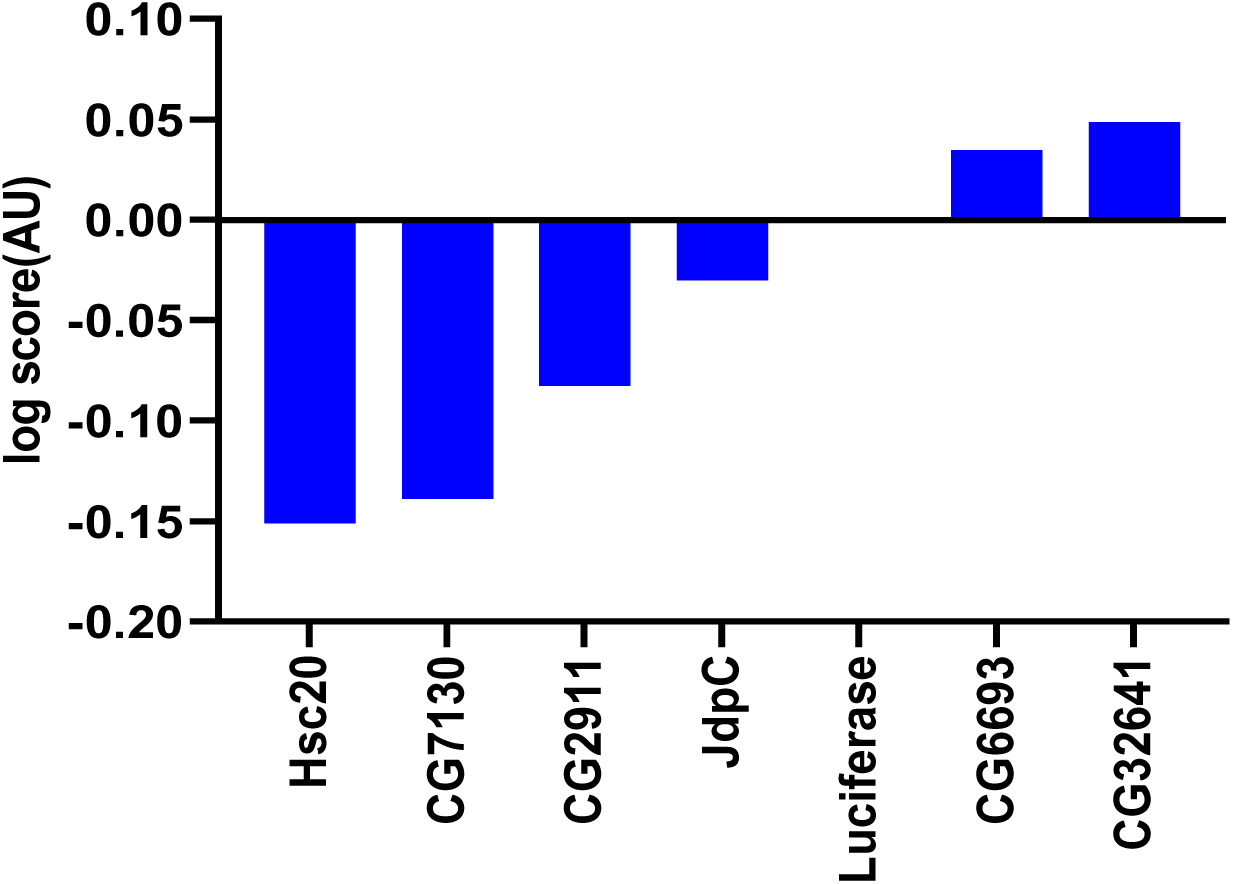
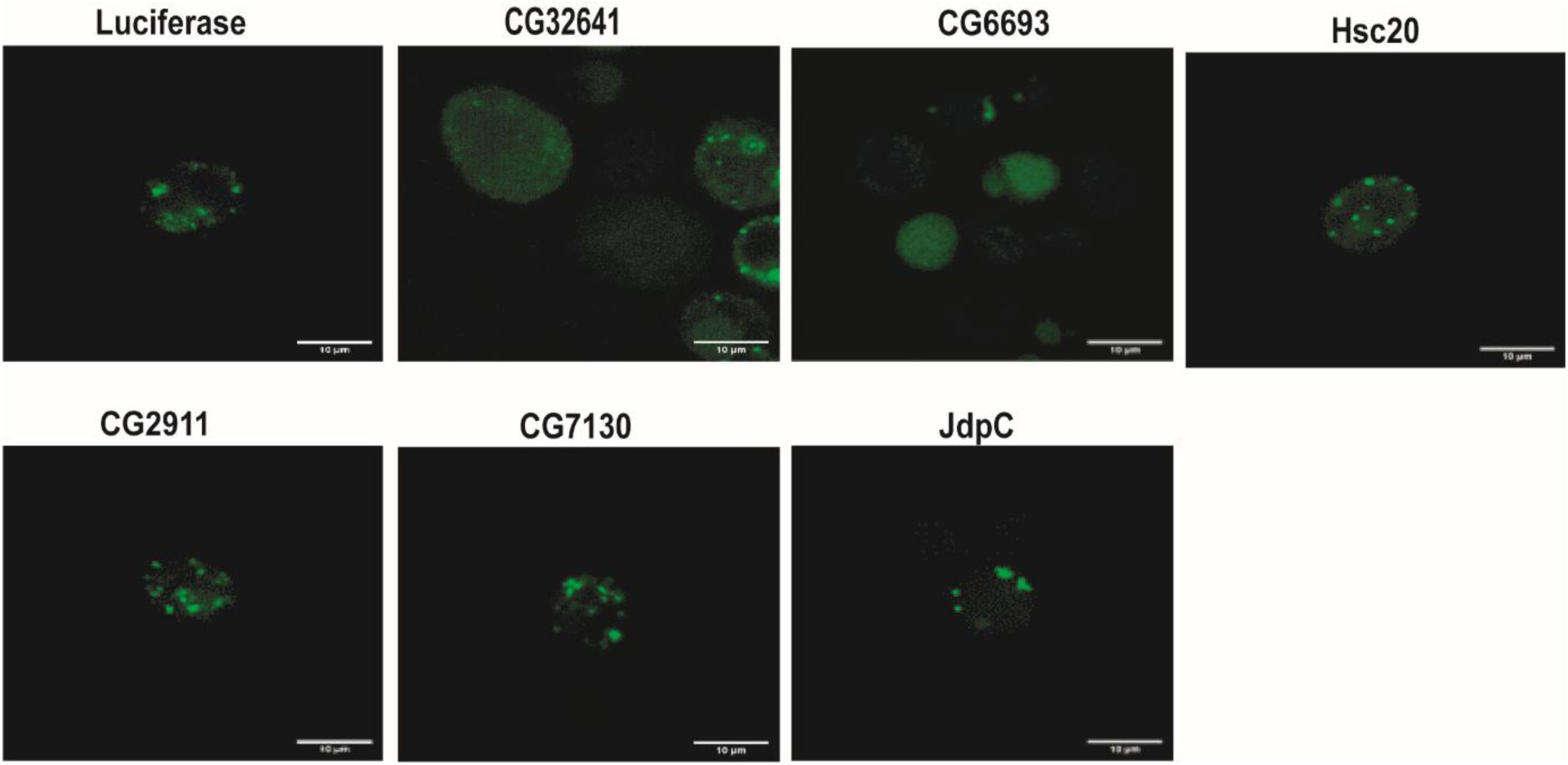
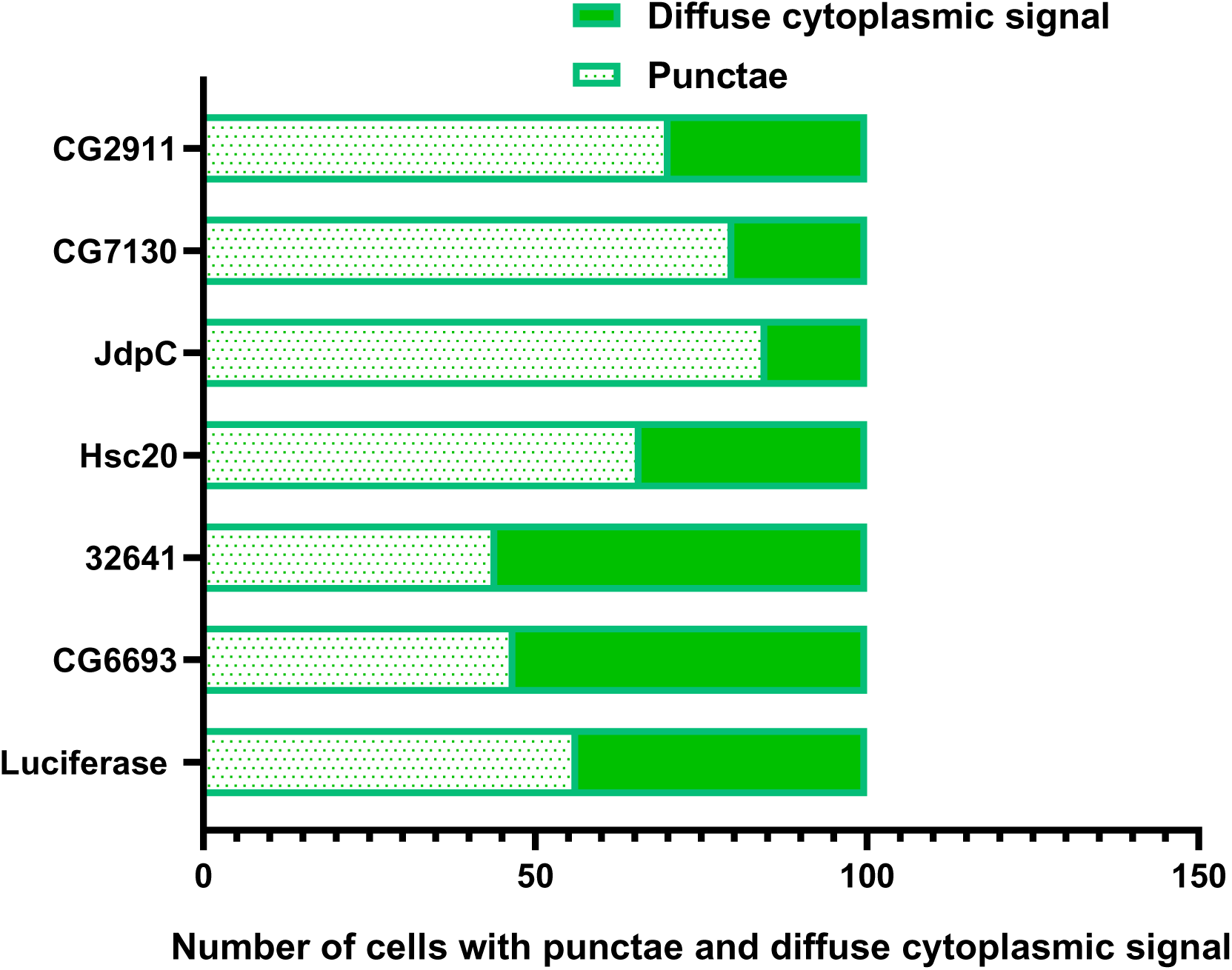
Effect of DnaJ domain chaperones on the growth of ALS mutant TDP-43: **a.** DnaJ domain chaperones transformed with ALS mutant TDP-43 were serially diluted and spotted onto SD-Trp-Ura and SGal-Trp-Ura plates. In this figure, SGal-Trp-Ura was shown as a test with TDP-43, and luciferase was used as a control. **b.** 48hrs growth curve of DnaJ domain chaperone transformed with TDP-43 mutant. Three independent biological replicates were used (n=3, P<0.001, paired t-test). The blue line indicates mutant TDP-43 with luciferase as control, and the red line indicates the DnaJ domain chaperones with the TDP-43 mutant. **c.** Enhancers and suppressors of TDP-43 mutant plotted, based on the area under the curve values obtained from the growth curves using GraphPad Prism software. The area under the growth curve values are converted into a logarithmic scale and normalized. Chaperones on the right-hand side are the suppressors, and those on the left-hand side are enhancers. **Fig. 1 Effect of DnaJ domain chaperones on protein aggregation phenotype of the TDP-43 mutant of ALS** **d.** Microscopic images showing protein aggregation and diffuse cytoplasm phenotype in the *Saccharomyces cerevisiae* model of ALS. TDP-43 aggregates are seen in the form of bright GFP punctae after induction with 20% Galactose. **e.** Two different morphologies exhibited by the different DnaJ domain chaperones with the TDP-43 mutant. 100-120 cells were counted for each chaperone. The average score was plotted using GraphPad prism. Table of total diffuse cytoplasmic signal and punctae.

**Table 2.** Shows the quantification of punctae and diffuse cytoplasmic signal of 7 DnaJ domain chaperones.

|  | Puncta | Diffuse cytoplasmic signal |
| --- | --- | --- |
| Luciferase | 60.10 | 39.90 |
| CG7872 | 40.22 | 59.70 |
| Mrj | 39.20 | 60.80 |
| Hsc70-4 | 40.00 | 60.00 |
| Hsc70-1 | 45.00 | 55.00 |
| CG2911 | 88.80 | 11.11 |
| CG2790 | 66.80 | 33.20 |
| CG8476 | 82.50 | 17.50 |

Chaperones that made the ALS mutants grow better in SGal-Trp-Ura plates than control, i.e., luciferase suppresses the effect of the ALS mutation, and are referred to as suppressors. Chaperones Mrj, Hsc70-4, and CG7872 are suppressors of TDP-43 toxicity (**Fig. 1a).** In the SD-Trp-Ura plate, they serve as control **(Supplementary Fig. S4a).** Chaperones CG5001 (previously published) (Deo et al., 2024), CG6693 and CG32641 grow better in SGal-Trp-Ura plates than luciferase. These chaperones are suppressors of FUS toxicity (**Fig. 1f).** SD-Trp-Ura plates shows no difference in growth **(Supplementary Fig. S5a).**

To quantify the growth effect, we did a 48-hour growth curve assay of DnaJ chaperones on TDP-43 and FUS mutants. TDP-43 and FUS transformants were inoculated in an SD-Trp-Ura medium for primary culture. The following day, O.D. was adjusted, and 20% galactose was added. Cultures were added in triplicate in a 96-well plate and placed in a Biotek Epoch spectrophotometer for 48 hrs at 30°C for occasional shaking. The data obtained from this experiment were processed in GraphPad Prism 8.0. Similar to the spot dilution assay, This 48-hour growth curve shows a growth difference compared to the luciferase.

Chaperones CG2911, CG2790, and CG8476 grows slower than control in a 48-hour growth curve. This indicates that these chaperones enhance the toxicity of mutant TDP-43. Chaperones Mrj, Hsc70, and CG7872 grow better than control, suggesting these chaperones rescue TDP-43 toxicity (**Fig. 1 b)**.

In FUS, chaperones Hsc20, CG2911, CG7130, and JDPC in presence of the FUS mutant grows slower compared to luciferase. This indicates that these chaperones enhance the toxicity of FUS. Co-expression of FUS with the chaperones CG6693, CG32641 and CG5001(Deo et al., 2024) have been shown to rescue the slow growth phenotype (**Fig. 1g).** In SD-Trp-Ura plates there is no difference **(Supplementary Fig. S5a).**

For the classification of the enhancers and suppressors of DnaJ domain chaperones in the 48-hour growth curve, we calculated the area under the curve based on the values of the 48-hour growth curve. The area under the curve values were normalized with luciferase. A graph was plotted in GraphPad Prism 8.0. Positive values of chaperones on the right-hand side are considered suppressors. The negative values of chaperones on the left-hand side are called enhancers (**Fig. 1c and 1h).**

### 2.3 Classification of weak and ineffective enhancers of *Drosophila* DnaJ domain chaperones

DnaJ domain chaperones of *Drosophila* were screened to see the effect on ALS mutant TDP-43 and FUS, after the yeast transformation of ALS mutants with DnaJ domain chaperones and co-chaperones. Initially, we checked by using a serial dilution assay. Positive clones were selected from the transformation and inoculated in SD-Trp-Ura (dextrose, YNB, and Tryptophan and Uracil lacking medium) (B. S. Johnson et al., 2008b). After 48 hrs, serially diluted cultures were spotted on SGal-Trp-Ura and SD-Trp-Ura plates. Plates with dextrose (SD) serve as a control **(Supplementary Fig. S4b and S5b)**, while galactose plates were observed for growth, as ALS plasmids have galactose-inducible promoters.

To quantify the effect of mutants on DnaJ domain chaperones, we did 48 hrs growth curve assay. ALS mutant TDP-43 and FUS with control luciferase and transformed chaperones were subjected to induced 20% galactose. These were grown at 30°C for 48 hrs, and the O. D_600nm_ with constant shaking, measured every 30 minutes. From 3 independent biological independent replicates, graphs were plotted in GraphPad Prism 8.0 software (**Fig. 2b and 2g). T**h**e** red line shows the mutants with transformed DnaJ domain chaperones, and the blue line shows the growth curve of ALS mutants with empty vector. (n=3, P<0.05, paired t-test).

The area under the curves is calculated for each chaperone, including luciferase converted into logarithmic values. Each value was further normalized with luciferase. A graph was plotted in GraphPad Prism 8.0 software. Chaperones that grow slower with ALS mutants were aligned to the right-hand side with a positive value were called suppressors, and the chaperones that were aligned to the left-hand side with a negative value are called enhancers (**Fig. 2c and 2h).** The classification of enhancers and suppressors of other chaperones is based on the values of the growth curve obtained from area under the curve.

In serial dilution assay and 48 hrs growth curve of TDP-43, chaperones CG12020, CG7394, CG4164, DnaJ60, CG3061, Hsp70Aa, CG32641, DnaJ1, JdpC, CG2887, Hsc20 and JdpA, grow less in comparison to the control. These are classified as weak enhancers. Other chaperones like CG14650, Tpr2, CG9828, CG7556, P58IPK, and Hsp70Bb grow equally to Luciferase and show no effect on the growth (n=3, P<0.05, paired t-test) shown in (**Fig. 2a).** In serial dilution assay and 48 hrs growth curve of FUS, chaperones CG9828, CG12020, Hsp70Aa, Hsp70Bb, CG11035, CG3061, DnaJ60, CG7133, CG14650, CG2887, CG10375 grow less compared to luciferase, are classified as weak enhancers, whereas chaperones CG7556, CG17187, CG43322, Csp, Sec63, CG11035, and Hsc70-1 show no effect on growth in the growth curve (**Fig. 2f).**

**Fig. 2.**
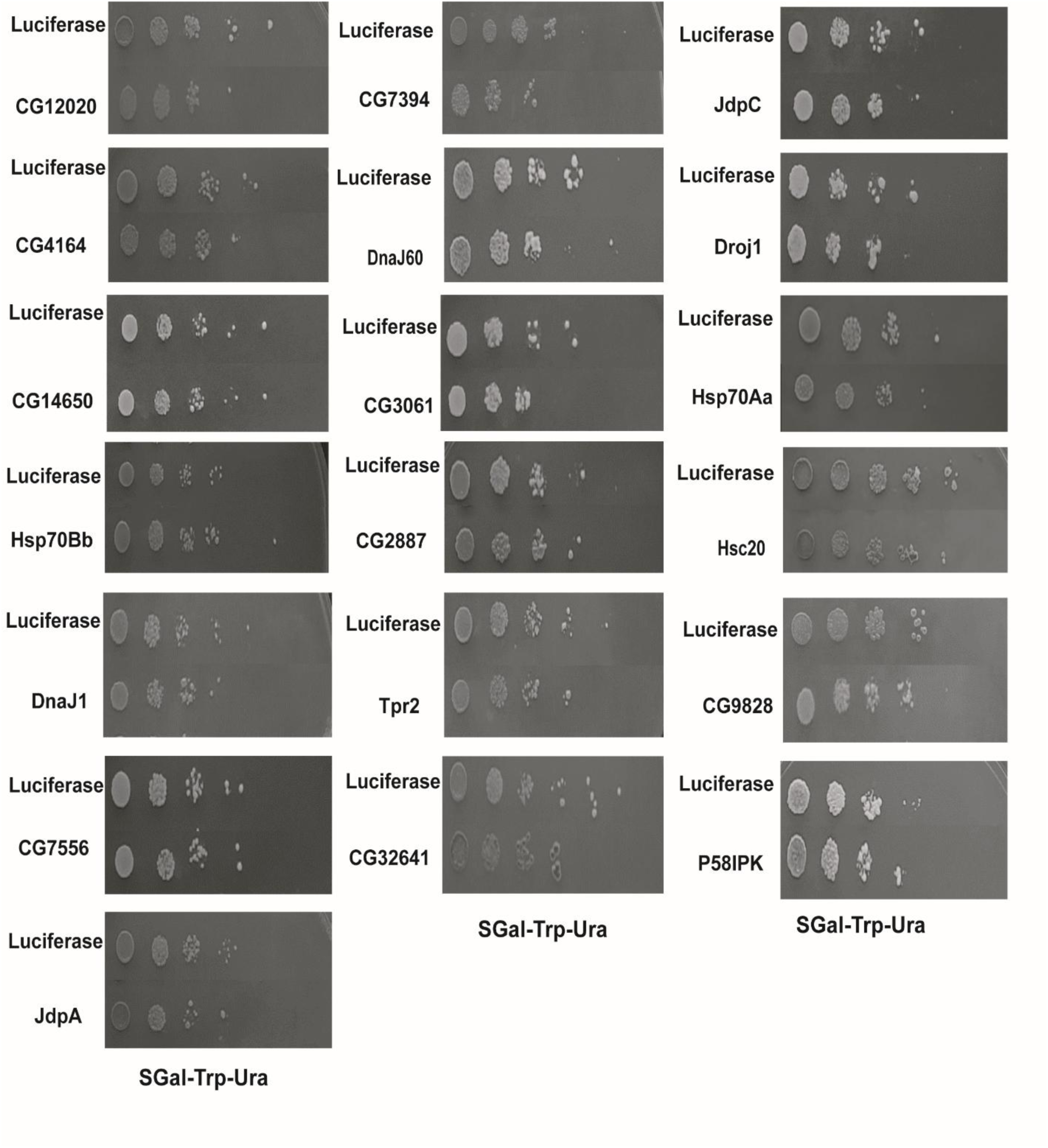
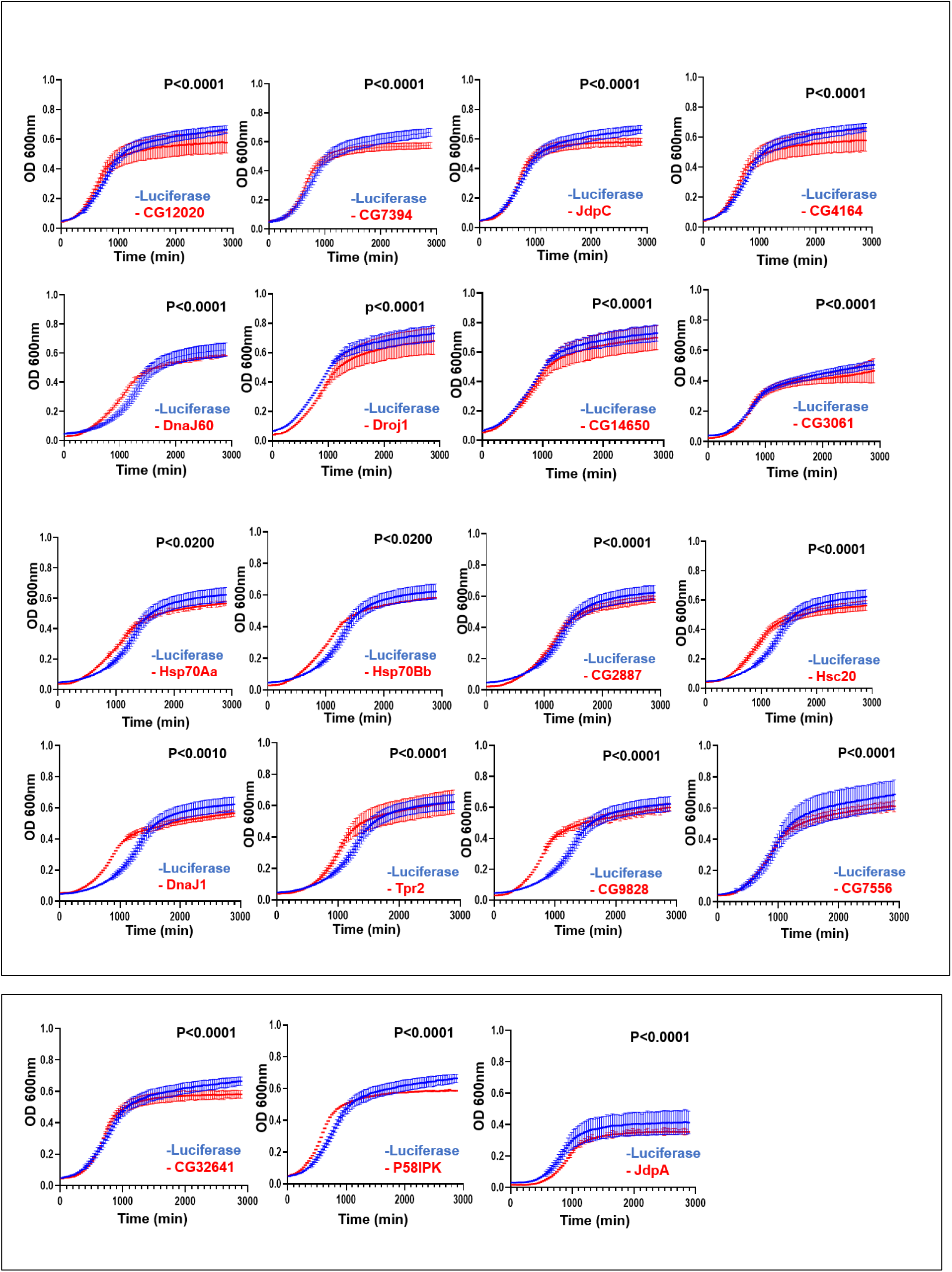
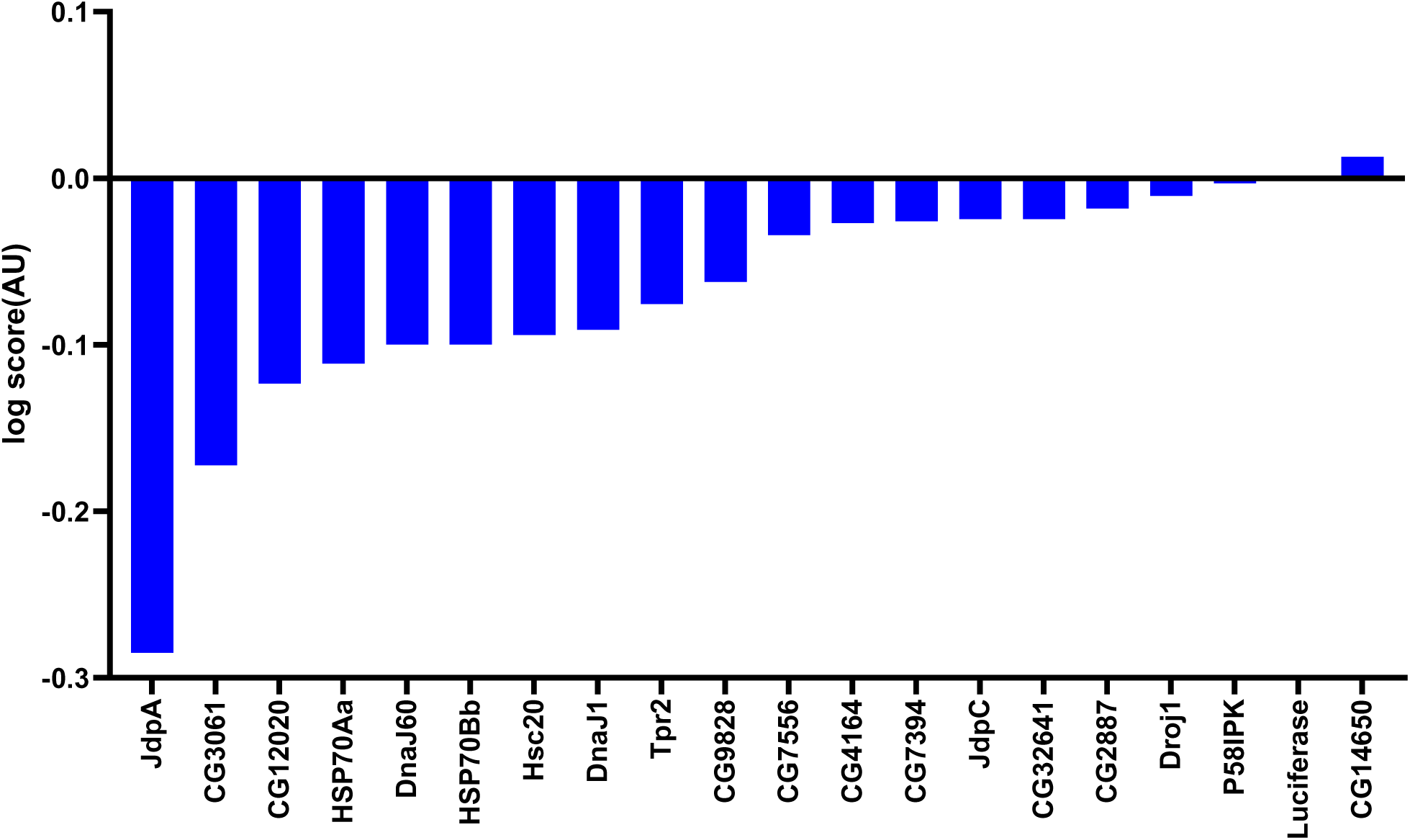
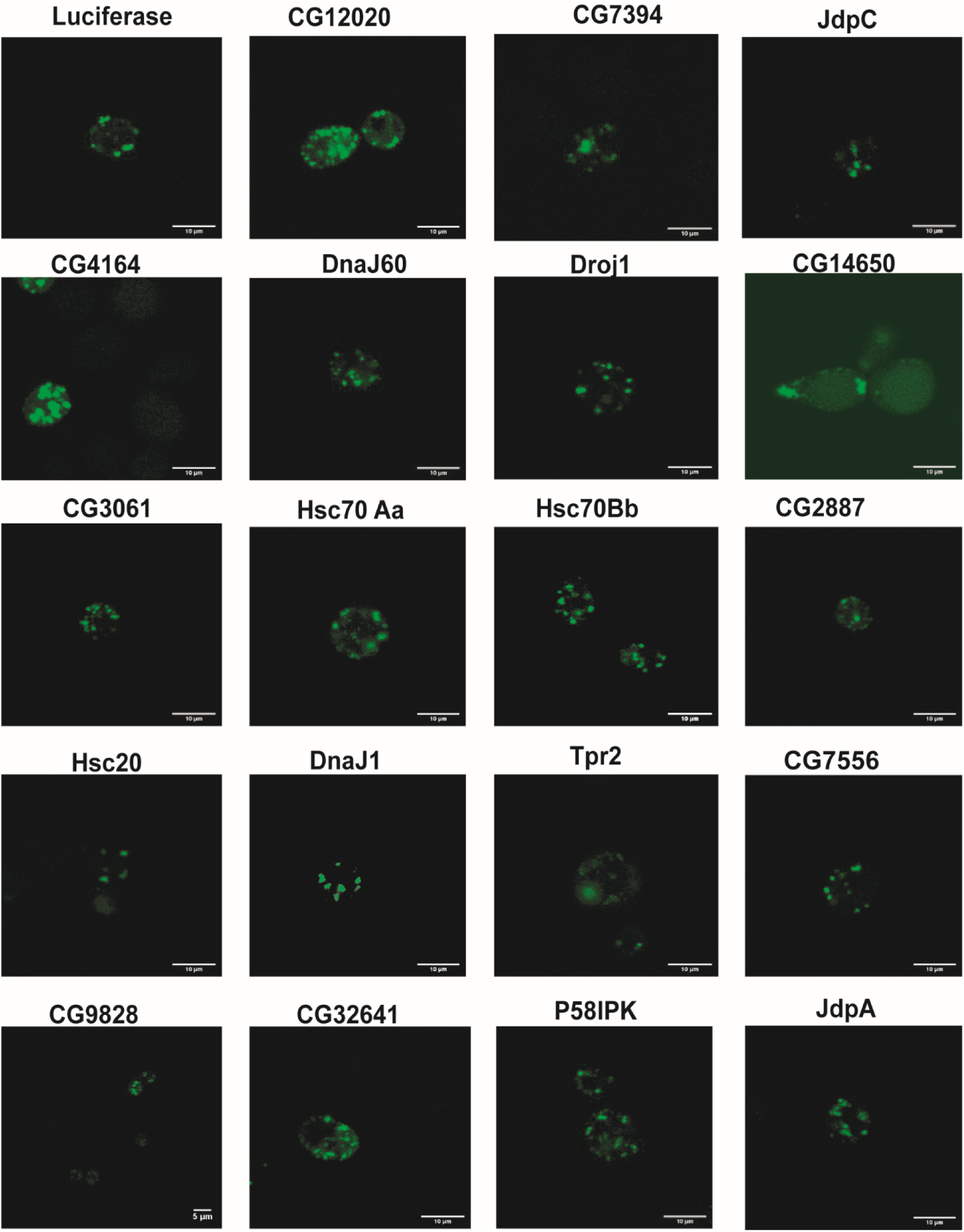
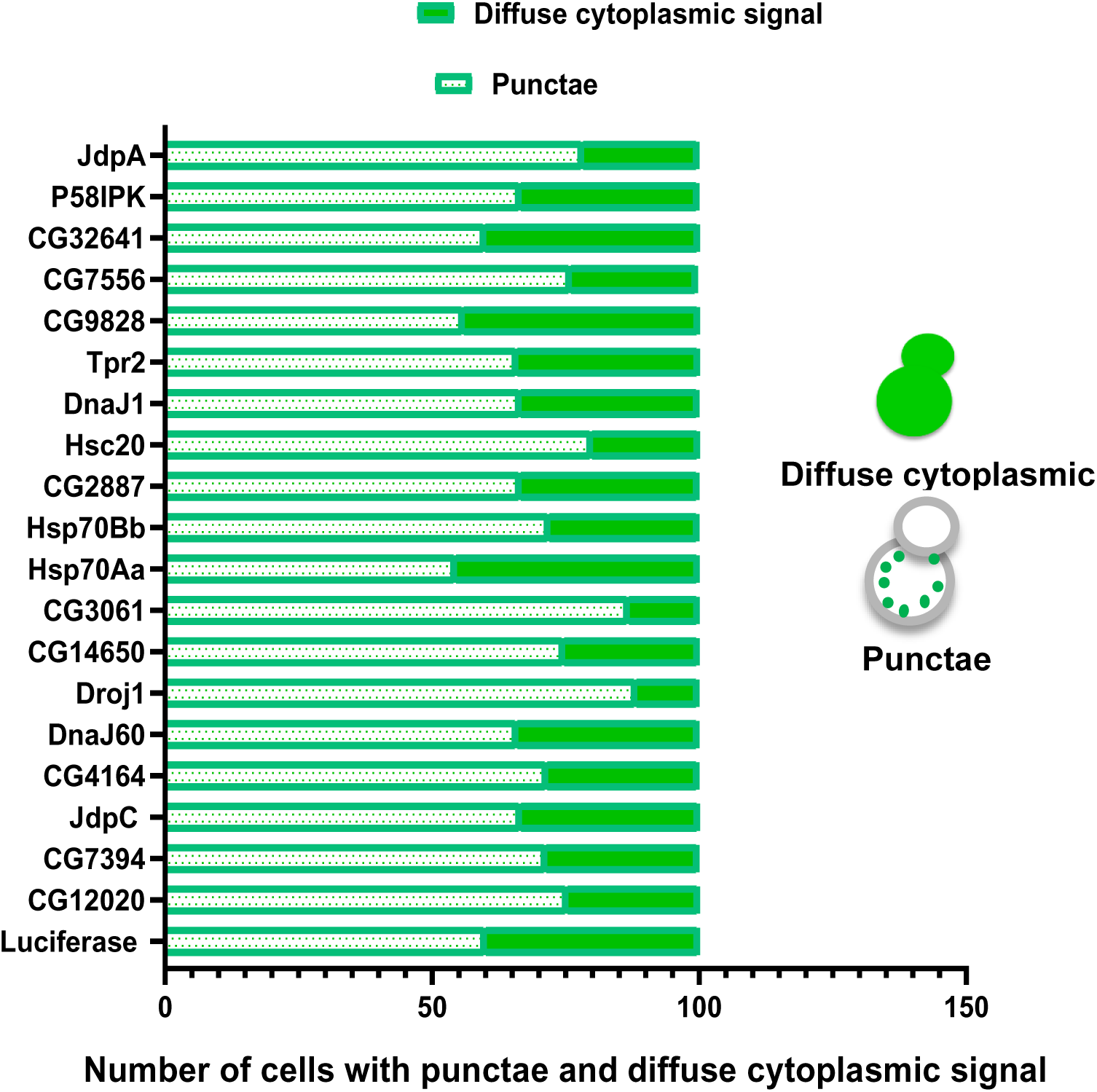
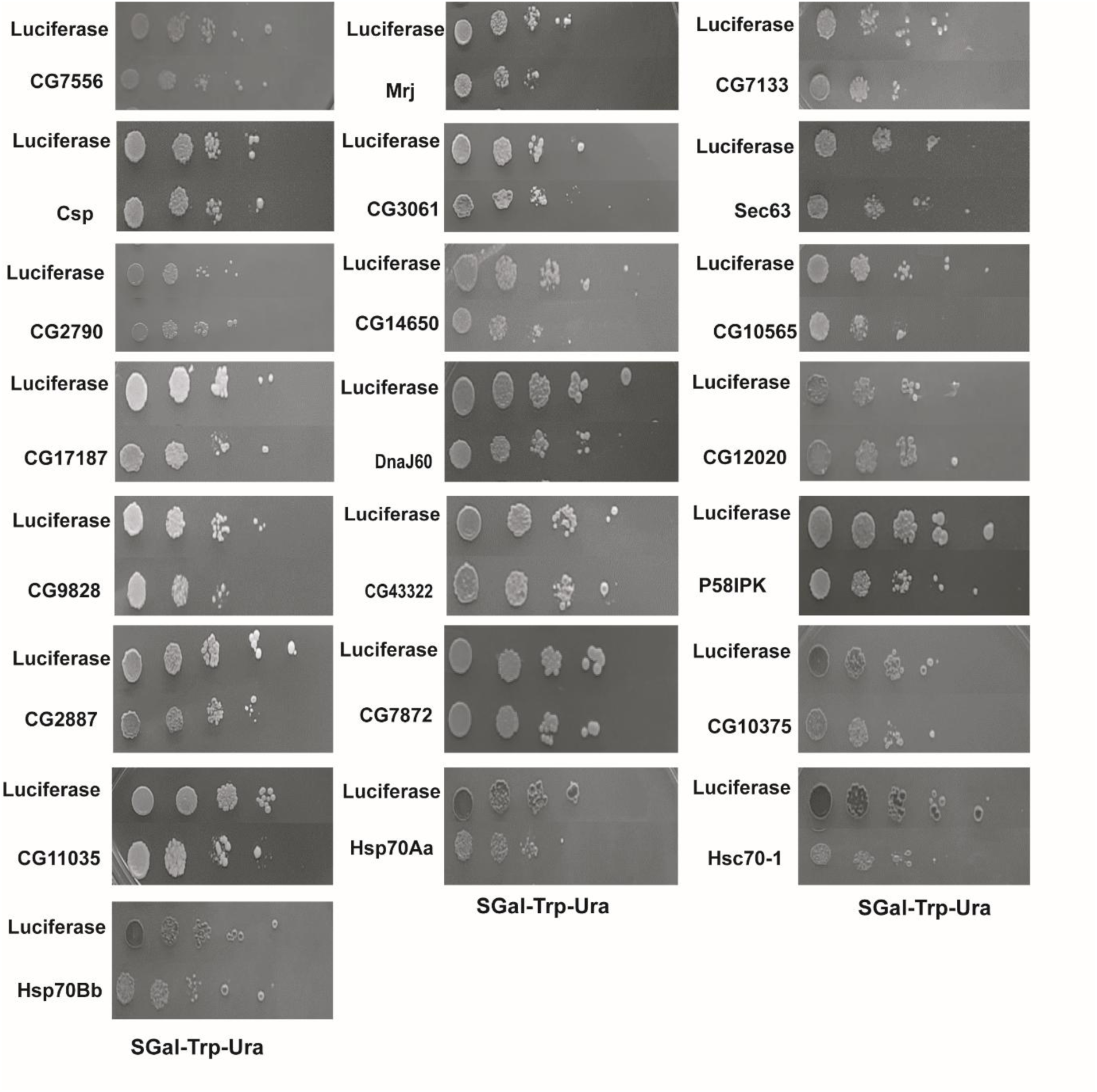
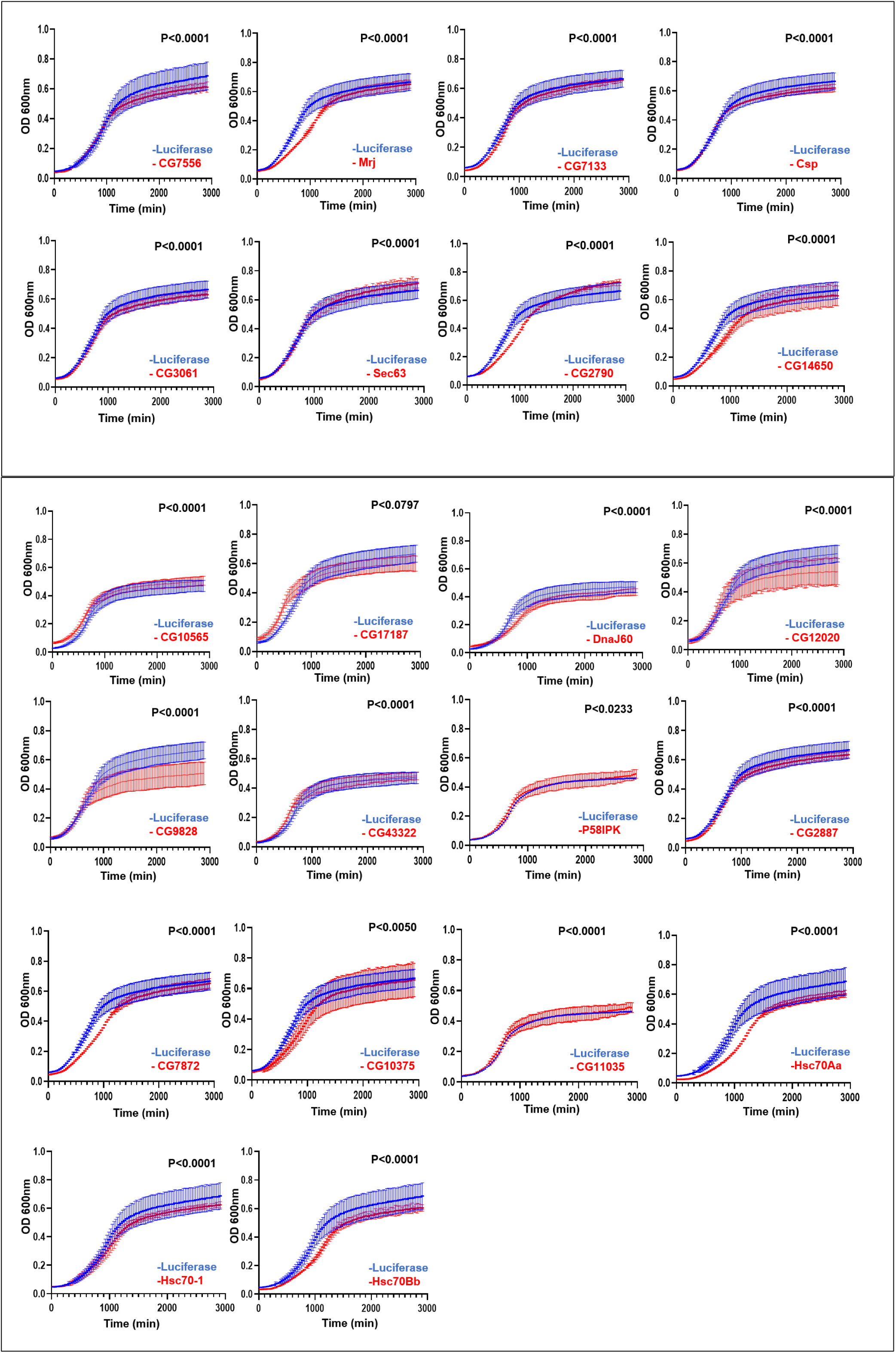
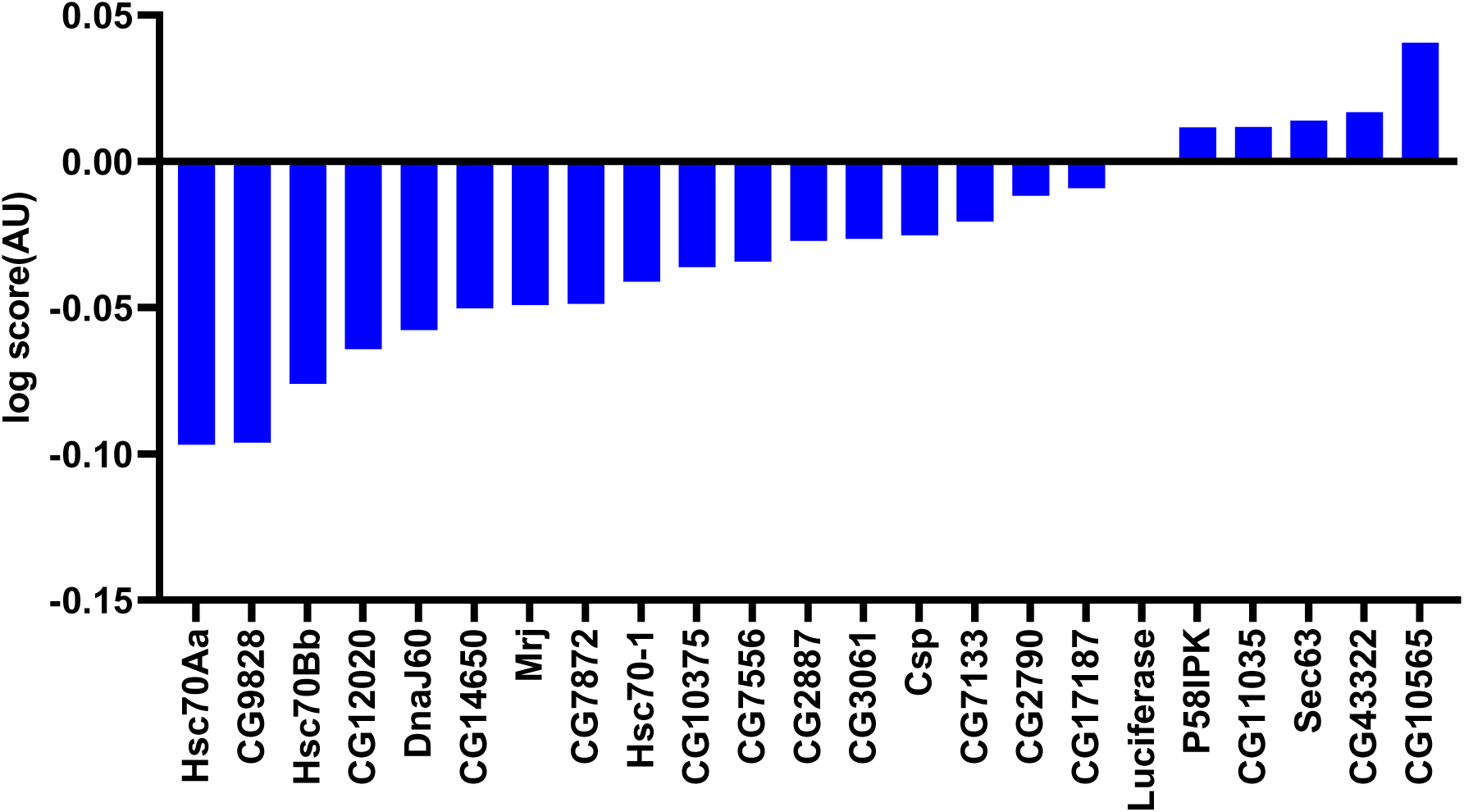
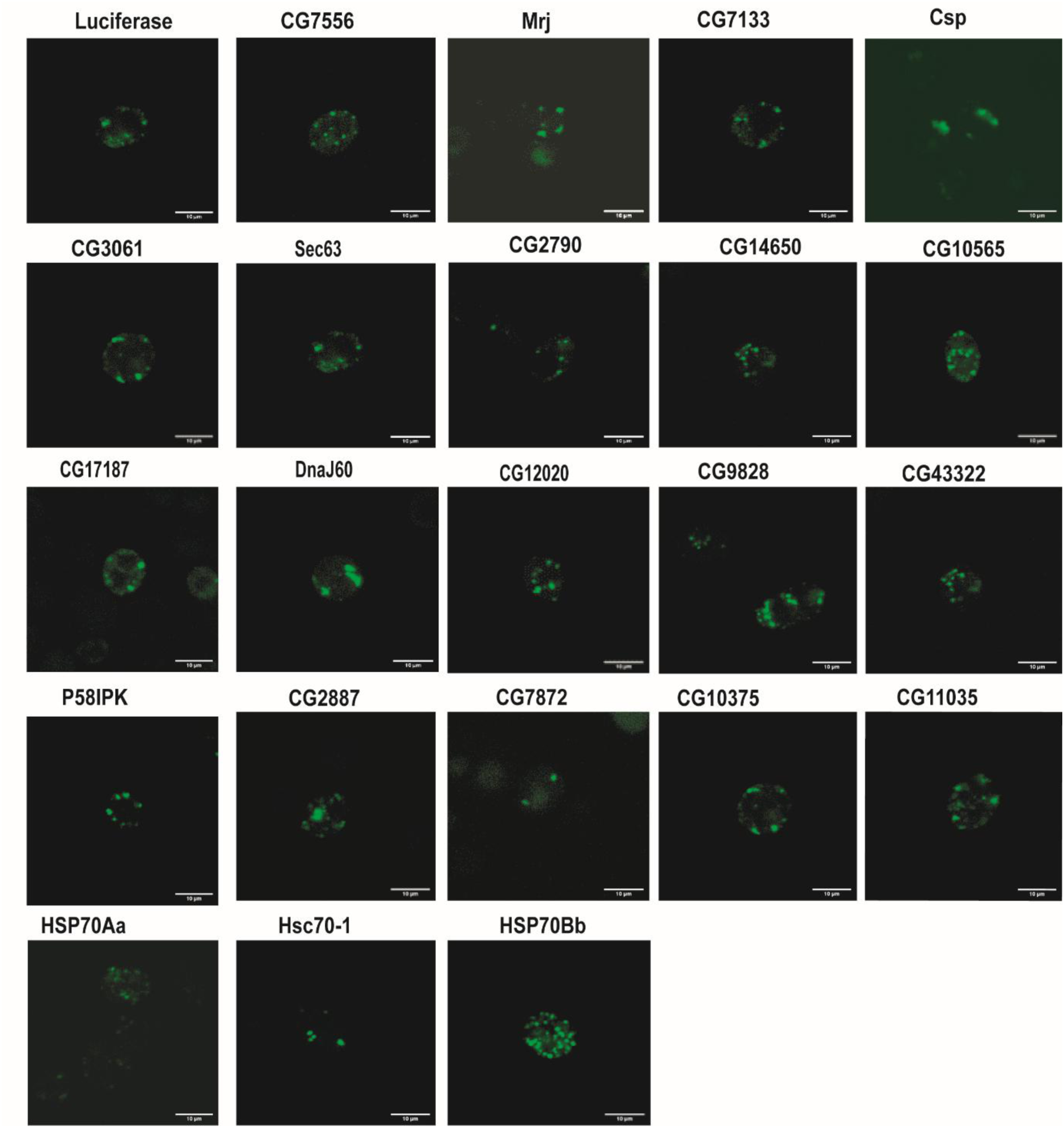
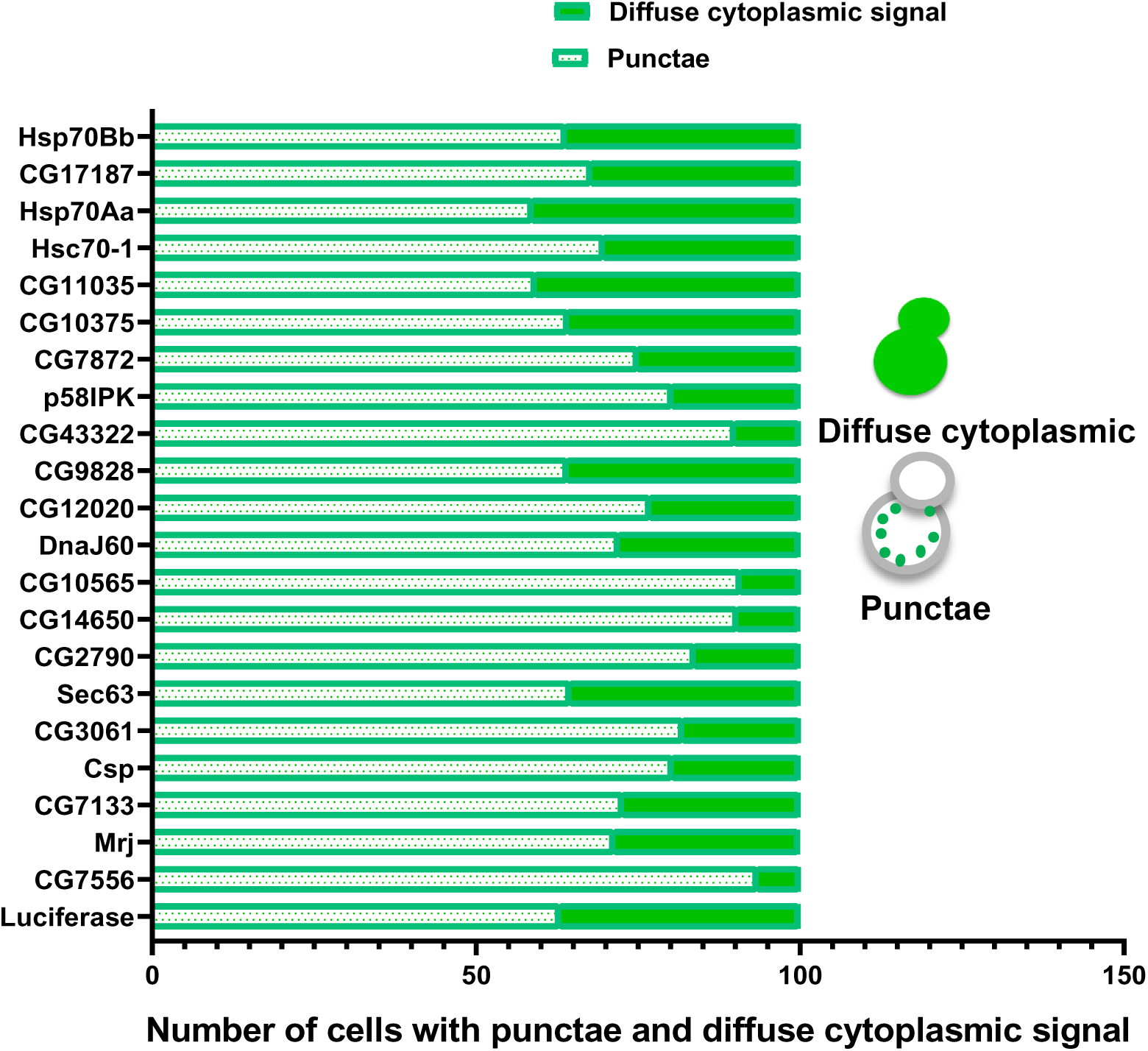
Effect of growth on ALS mutant due to chaperones (Serial dilution assay on Galactose medium) **a.** Representative image of serial dilution assay on SGal-Trp-Ura medium of DnaJ chaperones and co-chaperones mentioned in the study. **b.** Effect of chaperones on the growth of ALS mutant TDP-43 (48-hour growth curve) 48-hr growth curve of the remaining 19 DnaJ domain chaperones with Luciferase. Data shown in **Figure 2b** were obtained from three independent biological replicates (n=3, P<0.05, paired t-test) **c.** Enhancer and suppressor classification of the 19 DnaJ domain chaperones with TDP-43 mutant based on area under the curve values derived from GraphPad Prism. The area under the curve values of each chaperone were converted to a logarithmic scale and normalised. It is plotted via GraphPad Prism and the area under the curve values of the graph on the right-hand side are the suppressors, and those on the left-hand side are the enhancers. **d.** Effect of chaperones on protein aggregation phenotype of TDP-43 mutant. 19 different DnaJ chaperones and co-chaperones showing protein aggregation in the yeast model of ALS. Following induction with 20% galactose, microscopic images were taken. TDP-43 appeared as bright GFP punctae. **e.** Graph of 19 chaperones plotted in Graphpad Prism shown in **Figure 2c**. For each chaperone, 100-120 cells were counted, and the average was plotted. **g.** DnaJ domain chaperones transformed with ALS mutant FUS was serially diluted and spotted onto SD-Trp-Ura and SGal-Trp-Ura plates. In this figure, SGal-Trp-Ura was shown as a test with FUS luciferase as a control. **h.** 48-hour growth curve of DnaJ domain chaperone transformed with FUS mutant. Data from three independent biological replicates were collected (n=3, P<0.001, paired t-test). The blue line indicates mutant FUS with luciferase as control, and the red line indicates DnaJ domain chaperones with the FUS mutant. **i.** Enhancers and suppressors of FUS mutant based on the area under the curve values of obtained from the growth curves using GraphPad Prism software. The area under the growth curve values are converted into logarithmic scale and normalized. Chaperones on the right-hand side are the suppressors, and on the left-hand side, chaperones are enhancers **j.** Microscopic images showing protein aggregation and diffuse cytoplasm phenotype in the *Saccharomyces cerevisiae* model of ALS. FUS aggregates are seen in the form of bright GFP punctae after induction with 20% Galactose. **k.** Two different morphologies exhibited by the different DnaJ domain chaperones with the FUS mutant. Counted 100-120 cells for each chaperone average was plotted on GraphPad Prism. **Fig. 2 Effect of growth on ALS mutant FUS due to chaperones (Spot dilution assay on Galactose medium)** **f.** Representative image of serial dilution assay on SGal-Trp-Ura medium of DnaJ chaperones and co-chaperones with the FUS mutant mentioned in this study shows a weak enhancer, and some show no effect. **g.** 48-hour growth curve of DnaJ domain chaperone transformed with FUS mutant. Three biological replicates of data were collected (n=3, P<0.001, paired t-test). The blue line indicates mutant FUS with control Luciferase, and the red line indicates DnaJ domain chaperones with the FUS mutant described as a weak enhancer, and some show no effect on the mutant FUS **h.** Enhancer and suppressor classification of the remaining 21 DnaJ domain chaperones with FUS mutant, based on the area under the curve values derived from GraphPad Prism. The area under the curve values of each chaperone were converted to a logarithmic scale and normalized. It is plotted via GraphPad Prism and the area under the curve values graph on the right-hand side are the suppressors, and those on the left-hand side are the enhancers. **Fig 2. Effect of DnaJ domain chaperones on the protein aggregation phenotype of the FUS mutant of ALS.** **f.** Different DnaJ chaperones and co-chaperones are showing protein aggregation in the yeast model of ALS. following 20% galactose, microscopic images were taken protein aggregation was seen in the form of bright GFP punctae, indicating the weak enhancer. Some chaperones show both diffuse cytoplasmic signal and punctae phenotype as per the growth assay, and microscopic analysis indicates they are either enhancers or suppressors and do not show a strong effect on the mutant FUS. **g.** Graph of 21 chaperones plotted in Graphpad Prism is shown in Figure 2j. For each chaperone, 100-120 cells were counted, and the average was plotted.

**Table 3.** Shows the quantification of punctae and diffuse cytoplasmic signal of 19 DnaJ domain chaperones.

|  | Puncta | Diffuse cytoplasmic signal |
| --- | --- | --- |
| Luciferase | 60.10 | 39.90 |
| CG12020 | 75.38 | 24.62 |
| CG7394 | 71.40 | 28.50 |
| JdpC | 66.66 | 33.34 |
| CG4164 | 71.60 | 28.30 |
| DnaJ60 | 66.00 | 33.90 |
| Droj1 | 88.30 | 11.60 |
| CG14650 | 74.79 | 25.20 |
| CG3061 | 86.88 | 13.11 |
| Hsp70Aa | 54.54 | 45.45 |
| Hsp70Bb | 72.00 | 27.90 |
| CG2887 | 66.60 | 33.30 |
| Hsc20 | 80.00 | 20.00 |
| DnaJ1 | 66.60 | 33.30 |
| Tpr2 | 66.00 | 34.00 |
| CG9828 | 56.00 | 44.00 |
| CG7556 | 76.00 | 23.68 |
| CG32641 | 60.00 | 40.00 |
| P58IPK | 66.60 | 33.30 |
| JdpA | 78.30 | 21.60 |

**Table 4.**
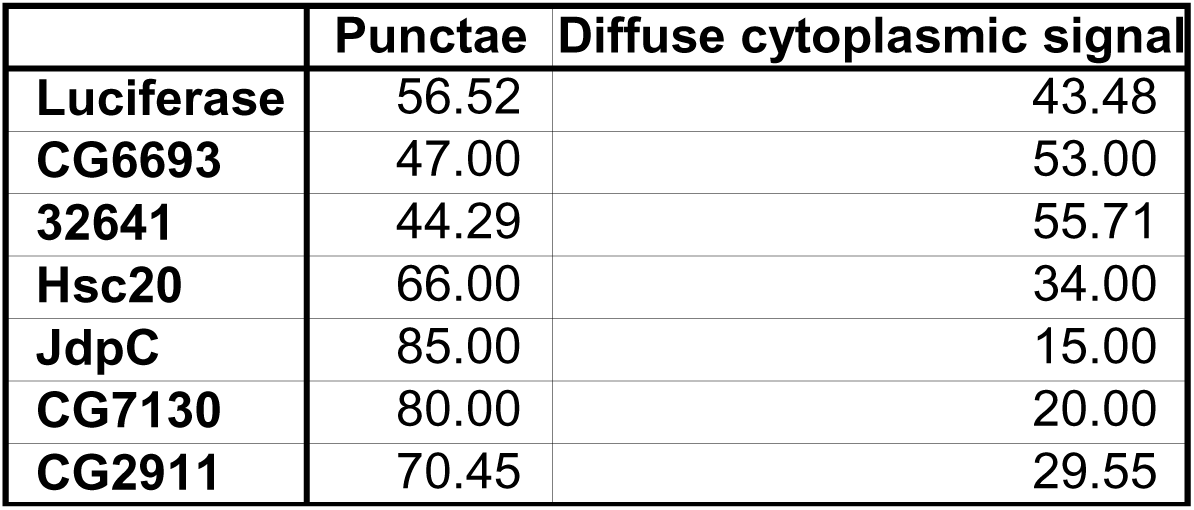
Shows the quantification of punctae and diffuse cytoplasmic signal of 6 DnaJ domain chaperones.

|  | Punctae | Diffuse cytoplasmic signal |
| --- | --- | --- |
| Luciferase | 56.52 | 43.48 |
| CG6693 | 47.00 | 53.00 |
| 32641 | 44.29 | 55.71 |
| Hsc20 | 66.00 | 34.00 |
| JdpC | 85.00 | 15.00 |
| CG7130 | 80.00 | 20.00 |
| CG2911 | 70.45 | 29.55 |

**Table 5.**
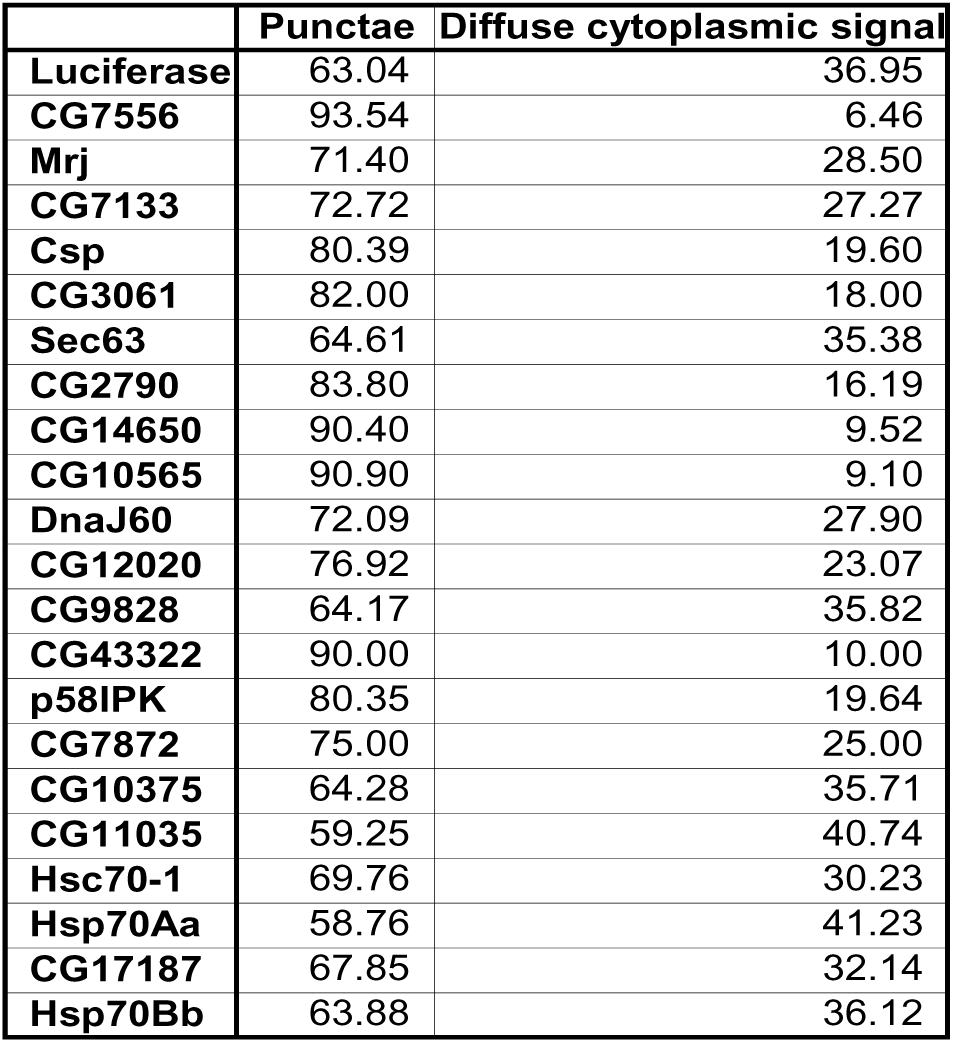
Shows the quantification of punctae and diffuse cytoplasmic signal of 21 DnaJ domain chaperones.

|  | Punctae | Diffuse cytoplasmic signal |
| --- | --- | --- |
| Luciferase | 63.04 | 36.95 |
| CG7556 | 93.54 | 6.46 |
| Mrj | 71.40 | 28.50 |
| CG7133 | 72.72 | 27.27 |
| Csp | 80.39 | 19.60 |
| CG3061 | 82.00 | 18.00 |
| Sec63 | 64.61 | 35.38 |
| CG2790 | 83.80 | 16.19 |
| CG14650 | 90.40 | 9.52 |
| CG10565 | 90.90 | 9.10 |
| DnaJ60 | 72.09 | 27.90 |
| CG12020 | 76.92 | 23.07 |
| CG9828 | 64.17 | 35.82 |
| CG43322 | 90.00 | 10.00 |
| p58IPK | 80.35 | 19.64 |
| CG7872 | 75.00 | 25.00 |
| CG10375 | 64.28 | 35.71 |
| CG11035 | 59.25 | 40.74 |
| Hsc70-1 | 69.76 | 30.23 |
| Hsp70Aa | 58.76 | 41.23 |
| CG17187 | 67.85 | 32.14 |
| Hsp70Bb | 63.88 | 36.12 |

The area under the curve of TDP-43, chaperones CG10565, CG12020, CG3061, DnaJ1, CG2911, CG7133, CG4164, DnaJ60, Droj1, and CG7394, JdpA, and JdpC are aligned to the left-hand side. These are classified as weak enhancers **(Fig. 2c)**. Whereas chaperones CG14650, Tpr2, CG9828, CG7556, P58IPK, and Hsp70Bb show no significant effect on growth. They are either aligned to the right-hand side or left-hand side and may be classified as intermediate.

The area under the curve of FUS, chaperones CG9828, CG12020, Hsp70Aa, Hsp70Bb, CG11035, CG3061, DnaJ60, CG7133, CG14650, CG2887, and CG10375 are aligned to the left-hand side and assigned a negative value. These are called weak enhancers **(Fig. 2h).**

Chaperones CG7556, CG17187, CG43322, Csp, Sec63, CG11035, P58IPK, and Hsc70-1 show no effect on growth in the 48-hour growth curve. They are aligned either to the left-hand side or the right-hand side. Therefore, they are classified as having no effect.

### 2.4 Enhancers and suppressors of ALS mutant TDP-43 and FUS based on protein aggregation in *S. cerevisiae*

Transformants of TDP-43 and FUS were induced with 20% galactose to express the ALS gene. Under confocal microscope, GFP expression is observed in yeast cells (B. S. Johnson et al., 2008b). TDP-43 and FUS showed puncta, a sign of protein aggregation, and the control showed diffuse cytoplasmic pattern with no protein aggregation inside the cell.

Around 100-120 cells were counted for ALS mutant TDP-43, FUS, and control independently. These were scored for the percentage of punctae and diffuse cytoplasmic GFP expression. Representative images are shown in (**Fig. 1d and 1i).** Based on confocal images, cells were classified into two types: cells with diffuse cytoplasmic expression and then with punctae. These were then compared to mutants with empty vector strains. The graph was plotted in GraphPad Prism. Chaperones, when overexpressed with mutant, have a higher expression of the cytoplasmic GFP diffusion than punctae. These chaperones are termed suppressors of the mutation. Those chaperones that showed greater number of punctae than diffuse cytoplasmic pattern was classified as enhancers. Chaperones Mrj, Hsc70-4, and CG7872 suppressed the effect of the mutation, and chaperones CG2911, CG2790, and CG8476 enhanced the mutation of TDP-43 in ALS (**Fig. 1e and Table 2)**. Chaperones CG5001(previously published) and CG6693 suppressed the effect of FUS toxicity, and chaperones Hsc20, CG2911, CG7130, and JdpC enhanced the FUS toxicity in the yeast cells (**Fig. 1i and Table 4**).

### 2.5 Classification of weak enhancers and no effect. Based on protein aggregation in TDP-43 and FUS

To check the protein aggregation of FUS and TDP-43 with the DnaJ domain chaperone, microscopy was done. Transformed cultures were inoculated in an SD-Trp-Ura medium (Dextrose, YNB, and Trp-Ura dropout). After 20% galactose induction, microscopy was done. Chaperones that show more of the diffuse cytoplasmic signal than punctae are called suppressors (CG10565, CG12020, CG3061, DnaJ1, CG2911, CG7133, CG4164, DnaJ60, Droj1). Strains with chaperones CG7394, JdpA, and JdpC on the other hand, show more punctae than diffuse cytoplasmic expression are the enhancers.

Microscopic images of TDP-43 with chaperones CG14650, Tpr2, CG9828, CG7556, P58IPK, and Hsp70Bb have more punctae compared to luciferase. These chaperones show mixed phenotype **(Fig. 2d).**

Microscopic images of FUS with chaperones CG9828, CG12020, Hsp70Aa, Hsp70Bb, CG11035, CG3061, DnaJ60, CG7133, CG14650, CG2887, and CG10375 show more of the punctae than control. Chaperones CG7556, CG17187, CG43322, Csp, Sec63, CG11035, P58IPK, and Hsc70-1 shows mixed phenotype (**Fig. 2i).**

For the quantitation of microscopic data, 100 cells were counted. Punctae and diffuse cytoplasmic morphology were counted for each transformant, and average values were plotted in GraphPad prism **(Fig. 2e and 2j).** Quantitation of both TDP-43 and FUS is given in Tables 3 and 5.

Chaperones of TDP-43, CG12020, CG7394, CG4164, DnaJ60, CG3061, Hsp70Aa, CG32641, DnaJ1, JdpC, CG2887, Hsc20 and JdpA have a higher percentage of punctae than diffuse cytoplasmic expression. Chaperones CG14650, Tpr2, CG9828, CG7556, P58IPK and Hsp70Bb show a mixed phenotype.

Chaperones of FUS, CG9828, CG12020, Hsp70Aa, Hsp70Bb, CG11035, CG3061, DnaJ60, CG7133, CG14650, CG2887, and CG10375 have a higher content of punctae compared to luciferase. Chaperones CG7556, CG17187, P58IPK, CG43322, Csp, Sec63, CG11035, CG10565 and Hsc70-1 shows a mixed phenotype.

## 3.0 Discussion

RNA-binding proteins TDP-43 and FUS plays a role in ALS pathogenesis. Both have some structural similarities, including the presence of a RNA-binding domain and a prion-like domain. They are both mutated in ALS (Gitler & Shorter, 2011a; Lagier-Tourenne et al., 2010). Although they have certain structural similarities, both genes contribute independently to ALS pathogenesis (Kwiatkowski et al., 2009b).

As per Arai and Newmann 2006, TDP-43 is localized in the nucleus and later spreads out to form cytoplasmic aggregates in ALS and FTLD patients (Arai et al., 2006; Neumann et al., 2006b).

Yeast cells do not have a true TDP-43 homolog but have RNA-binding proteins of similar architectural domain. TDP-43 might aggregate and sequester these proteins, which will offset their normal function [(Gitler & Shorter, 2011b).

In Parkinson’s, Huntington’s, Alzheimer’s, and prion-related studies, the respective protein forms, intracellular aggregates, and toxicity in yeast cells. Similarly, TDP-43 also forms cytoplasmic aggregates and can be added to the list of human misfolding diseases of proteins (Gitler, 2008).

FUS also forms cytoplasmic aggregates in neurons after a missense mutation (Vance, Rogelj, Hortobágyi, et al., 2009). FUS is normally located in the nucleus. After mutation, FUS forms pathogenic aggregates in motor neurons (Ju et al., 2011)

*In vitro studies* from yeast show TDP-43 and FUS tends to form cytoplasmic aggregates (Elden et al., 2010b; B. S. Johnson et al., 2008b, 2009). In animals and cellular models ATX2 gene is found to contribute to TDP-43 toxicity and ALS pathogenesis. It is a polyglutamine protein, and the human orthologue of yeast found mutated in spinocerebellar ataxia type 2 (Lorenzetti et al., 1997). Furthermore, it was observed that TDP-43 and ATX2 are associated in a complex (Elden et al., 2010b). In yeast, the pure form of the TDP-43 gene *in vitro* can form multiple aggregates *in vivo*. Additionally, a pure TDP-43 aggregates assay aids in identifying small molecules and cellular factors capable of impeding or reversing TDP-43 aggregates (B. S. Johnson et al., 2009). After dissecting the TDP-43 and expressing it in yeast, it was noticed that the C-terminal and RNA recognition motif contribute to TDP-43 toxicity (B. S. Johnson et al., 2008a).

Yeast is a genetically tractable model, and it is possible to reiterate multiple features of TDP-43 pathology in yeast (Gitler, 2008).This includes screening of multiple genomes for detecting modifications of protein aggregation and their toxicity. This is one approach that can be used for effective diagnosis and treatment (B. S. Johnson et al., 2008a).

In our observations, TDP-43 and FUS mutants showed slow growth in yeast cells after serial dilution, indicating cellular stress [(B. S. Johnson et al., 2008b; Sun et al., 2011). After 24 hours of galactose induction, these mutants show aggregates in the cell under the microscope.

Screening of these mutants with 38 DnaJ domain chaperones showed that some of the chaperones enhanced and others reduced the toxicity of the mutants in ALS. We labelled them as enhancers and suppressors of TDP-43 and FUS respectively.

By screening *Drosophila* DnaJ domain chaperones, we found that chaperones CG7872, Mrj, and Hsc70-4 can rescue the toxicity of mutant TDP-43. HSc70-1 slightly rescues the toxic phenotype of TDP-43 to a great extent. Hsc70-4. CG32641 and CG6693 suppresses the toxicity of mutant FUS. Previous reports show that, CG5001 rescues both the mutants TDP-43 and FUS (Deo et al., 2024). Human homologues of Hsp70, chaperones DNAJAs, DNAJBs, and DNAJCc chaperones rescue the toxicity of TDP-43 in ALS (Barbieri et al., 2025).

From our studies we see, chaperone CG2911 acts as a common enhancer for both TDP-43 and FUS.

Chaperones CG7872, Hsc70-4, and Mrj demonstrate a more significant growth effect in mutant TDP-43. In the serial dilution assay and the growth curve, they grow comparatively better than the control. In microscopy, they exhibit a higher prevalence of diffuse cytoplasm morphology than punctae; hence, we refer to them as suppressors of TDP-43.

DnaJ chaperone Mrj is present ubiquitously in the cytosol. Mrj’s human orthologue, DNAJB6, is known for its potent anti-aggregation effect in mouse models (Kakkar et al., 2016). A mutation in the human DNAJB6 (Mrj) causes frontotemporal dementia and limb-girdle muscular dystrophy type 1 case of the patient’s brain sample (Yabe et al., 2014). It was also found that DNAJB6 inhibits polyQ aggregation and increases the life span in polyQ mice (Kakkar et al., 2016). *Drosophila* Mrj helps Orb2 to reduce its prion conversion and also helps reduce Htt aggregates (Desai et al., 2024). DNAJC25 is an ortholog of CG7872, belonging to the HSP40 family C, and plays a vital role in various human diseases, including neurodegenerative disorders (LIU et al., 2012b). Co-chaperone Hsc70 influences TDP-43 toxicity in ALS patients (Arosio et al., 2020)

Hsc70-4 is the human orthologue of HPSA8 and has been found to control multiple studies of autophagy (Braell et al., 1984; Stricher et al., 2013). Previously, it was also found to prevent necroptosis initiation and bind to reverse RIP3 amyloid in mammalian cell culture(Wu et al., 2023)

In the FUS mutants, chaperones CG32641 and CG6693 come up as suppressors as they showed better growth than luciferase in both serial dilution and growth curve. Microscopic quantitation shows more of the diffuse cytoplasmic morphology and fewer punctae in the cell. Therefore, they are referred to as suppressors of FUS toxicity.

Reports show show that DNAJC9 is the human orthologue of CG6693 and is associated with cervical cancer. It is probably upregulated by reducing or protecting against apoptosis in the cancer cell line (Incekara & Acun, 2023). Studies in zebrafish stated that DNAJC9 acts in the downstream regulating pathway of the tumor suppressor gene p53 (Mandriani et al., 2016).

Also, CG32641 has a DNAJB9 human orthologue and is known for its ER-associated degradation machinery (Lai et al., 2012). In the mouse model, it was found that it rescues cystic fibrosis-causing gene Δ F508, and also directly interacts with CFTR (Huang et al., 2019).

From the studies of Drosophila DnaJ domain chaperones, we also discovered certain enhancers that exacerbate the toxic phenotype, specifically slow growth and aggregation in both mutant FUS and TDP-43. Chaperones CG2911, CG2790, and CG8476 enhance TDP-43 toxicity and have also been shown to be toxic in multiple studies. DNAJC21 is the human orthologue of CG2790, and its mutation is associated with bone marrow failure and cancer (Tummala et al., 2016). DNAJB9, is the human orthologue of CG8476, whose normal function involves protein folding. It is predominantly active in the endoplasmic reticulum and is found in the miR-32 gene in myeloid acute leukaemia (Thomson & Dinger, 2016).

Whereas, chaperones Hsc20, CG2911, CG7130, and JDPC are enhancers of mutant FUS. They also have a role in other disorders like Hsc20 which is also a mitochondrial chaperone, its human orthologue is HSCB; Mutation of Hsc20 in *Drosophila* larvae causes reduced iron-sulphur cluster enzyme activity (Uhrigshardt et al., 2013). JdpC has a human orthologue to DNAJC12, and in whole-exome sequencing, it is found responsible for hyperphenylalaninemia and global developmental delay in some children (Veenma et al., 2018).

CG2911 acts as an enhancer for in both TDP-43 and FUS. Its human orthologue is DNAJC24, which recently was reported to be involved in hepatocellular carcinoma (D. Liu et al., 2024). In our studies, it also shows toxicity when overexpressed in both mutants of TDP-43 and FUS.

While screening in the TDP-43 background we found that chaperones CG12020, CG7394, CG4164, DnaJ60, CG3061, Hsp70Aa, CG32641, DnaJ1, JdpC, CG2887, Hsc20 and JdpA are weak enhancers. This can be attributed to the slow growth phenotype of these chaperone in comparison to the mutant. In image analysis, accordingly, we labelled them as weak enhancers **[Table.1 (Fig. 2a, 2b**, **2c, and 2e)]**.

In the FUS strains, chaperones CG9828, CG12020, Hsp70Aa, Hsp70Bb, CG11035, CG3061, DnaJ60, CG7133, CG14650, CG2887, and CG10375 are weak enhancers. Their growth and spot dilution assay show slower growth compared to luciferase. Also, in microscopy, they show more puncta than luciferase. In the area under the curve graph, they are aligned on the left-hand side, and in microscopic quantitation, they show more puncta. Therefore, they are classified as the weak enhancers of FUS toxicity **[Table.1** (**Fig. 2f, 2g, 2h, and 2i)].**

Some chaperones do not show any significant change in the toxicity of mutant TDP-43 example, CG14650, Tpr2, CG9828, CG7556, P58IPK, and Hsp70Bb. Serial dilution and growth curve results show equal growth. It is difficult to conclude whether they are enhancers or suppressors.

Similarly, in FUS mutants some chaperones, CG7556, CG17187, P58IPK, CG43322, Csp, Sec63, CG11035, CG10565, and Hsc70-1 do not show significant effects in spot dilution and growth curve. From these data, it is difficult to conclude whether these chaperones have any effect on FUS toxicity.

In both the mutant TDP-43 and FUS, Hsc70Aa and Hsc70Bb chaperone have no effect or weak enhancer.

In our study, we used *Drosophila* DnaJ domain chaperones in an yeast background to look for potential suppressors and enhancers in both TDP-43 and FUS. Also, we find suppressors and enhancers that were previously unknown in ALS.

Based on the *Drosophila* DnaJ domain chaperone screening with FUS and TDP-43 mutants, we find that chaperone CG7872 reduces the toxicity of TDP-43 in yeast cells, which is previously unknown. There are many studies on Mrj, which show that it reduces the toxic aggregation in Huntington’s and Parkinson’s disease (Aprile et al., 2017; Desai et al., 2024; Kakkar et al., 2016). Based on our findings, we think Mrj possibly reduces the toxicity in ALS in a yeast model. Co-chaperone Hsc70 is found to reduce the toxic TDP-43 aggregation in sporadic ALS patients (Arosio et al., 2020).

We also find that chaperones CG32641 and CG6693 rescus the toxicity of FUS in yeast cells, which was not reported previously.

The hypothetical model in **Fig. 3** shows the mechanism of action.

**Fig. 3.** Hypothetical model. This schematic diagram is the overall hypothesis of our research, in which the ALS mutant genes of TDP-43 and FUS, when mutated, form misfolded proteins, resulting in an aggregation of protein within the cell. by interacting with Hsp40 chaperones, i.e, Drosophila DnaJ domain chaperones with yeast cells, they either enhance the toxicity or rescue the toxicity of these two mutants. Toxic phenotype in yeast represents puncta in the cytoplasm, and rescue represents cytoplasmic diffusion.

In this study, we have used a cloned library of 38 DnaJ domain chaperones of flies and expressed them in two ALS mutants, TDP-43 and FUS. This will help shed some light on the differences and similarities in the pathways regulated by TDP-43 and FUS in controlling the pathogenesis of ALS.

## 4.0 Material and Methods

### 4.1 Cloning of DnaJ domain chaperones and co-chaperones (Hsp40, Hsc70, and Hsp70) library in yeast

*Drosophila* Hsp40 and Hsp70 proteins were cloned by the Topo-D-Entr cloning kit (Invitrogen) into the Topo-D-Entr vector. This was done post-PCR amplification using gene-specific primers. Constructs were confirmed by sequencing and were transferred to the destination vector pUASgHA for Hsp40 (Bischof et al., 2013; Joag et al., 2020; Warrick et al., 1999) and pUASt-ccdB-FLAG (pTWF) for Hsp70.

For expression of Hsp40 and Hsp70 in yeast cells, Topo-D-Entr clones were cloned in the destination vector pAG424GALccdB-HA with a 3X HA tag at the C-terminal (Hageman et al., 2011). The protein expression of each isoform was confirmed by immunoblotting using either an anti-HA antibody or an anti-FLAG antibody accordingly (Desai et al., 2024).

### 4.2 Construction of yeast overexpression of DnaJ chaperones and co-chaperones in yeast TDP-43 and FUS strain backgrounds

#### 4.2.1 Bacterial Transformation

For the bacterial transformation, DH5αCells were made competent by using the calcium chloride method. 200μl competent cells were taken, and 1 μl plasmid DNA of ALS plasmids pRS426 Gal-TDP-43-GFP and pAG416 Gal-FUS-YFP were added. These were incubated on ice for 30 minutes. Then, heat shock was given for 2 minutes at 42°C, and it was kept on ice for 1 minute. Luria Broth was added to it, and incubated at 37°C for 30 to 45 minutes. The cells were centrifuged at 8000 rpm for 3 minutesat room temperature. Then, the supernatant was discarded, and the cells were resuspended in 200 μl of Luria broth. They were plated with glass beads on LB amp plates, and finally, the plates were incubated overnight (maximum 18 hours) at 37°C. These plasmids were deposited by Aaron Gilter lab, numbered 27467 and 29592, respectively(B. S. Johnson et al., 2008a; Sun et al., 2011).

#### 4.2.2 Plasmid Isolation

A single colony of transformed bacteria was inoculated in 5ml LB broth, and 5µl of Ampicillin was added and kept in a 37°C shaker overnight. The culture was centrifuged at 10,000 rpm for 5 minutes, and of GET was added. The culture was incubated on ice for 2 minutes. 200µl of alkaline SDS was added to it. The mixture was incubated on ice for 2 minutes, and 150µl acidic KoAc and incubated again on ice for 5 minutes. The whole culture was centrifuged at 14,000 rpm for 5 minutes at 4°C. The supernatant was collected, and 400µl of phenol: chloroform solution was added. It was centrifuged at 14,000 rpm for 5 minutes at 4°C. The aqueous phase was collected, and 1 mL of 100% ethanol was added to it. This was kept at -20°c for 15 minutes for precipitation and centrifuged at 14,000 rpm for 10 minutes at 4°C. The supernatant was decanted, and the pellet was washed with 500µl of 70% ethanol by centrifuging it at 14,000 rpm for 5 minutes at 4°C. The supernatant was discarded, and the pellet was air-dried. Once dried, the pellet was resuspended in 50µl of TE buffer. Finally, the plasmid was stored at - 20°C.

#### 4.2.3 Yeast Transformation

Yeast strain *Saccharomyces cerevisiae* (w303a, MATa, ade2, his3, leu2, trp1, ura3) cells were grown in YPD (yeast extract, peptone, and dextrose) medium, i.e., Primary culture, and allowed to grow overnight at 30°C, till mid log phase. After spinning down the cells at 5000 rpm for 5 minutes at room temperature, the pellet was resuspended in autoclaved distilled water to wash off residual media. The pellet was resuspended completely after spinning down in 100mM LiOAc, from which aliquots were made as per the number of transformations to be performed. Cells were centrifuged, and to each vial, 50% Polyethylene Glycol, 1 M Lithium acetate, ss DNA, and plasmid DNA were added (Ito et al., 1983). The cell suspension was vortexed to ensure resuspension and subjected to a heat shock at 42°C for 45 minutes in a water bath. The cell suspension was centrifuged at 5000 rpm for 5 min at room temperature, and autoclaved distilled water was used to resuspend the pellet.

The plasmids TDP-43 (pRS426 Gal-TDP-43-GFP) and FUS (pAG416 Gal-FUS-YFP) are galactose inducible and have an Uracil marker and are placed on SD-Ura, having a mixture of Sucrose, YNB, and amino acid lacking mixture Uracil. The media used were from Clontech Laboratories. USA. The plates were incubated at 30°C for 3-4 days (Gietz & Schiestl, 2007). In case of the DnaJ domain chaperone, double transformation was done with ALS mutants. These chaperones have a Trp marker. Transformation was done on SD-Trp-Ura (YNB, dextrose, agar, and amino acid mixture lacking Tryptophan and Uracil) medium.

#### 4.2.3 Serial dilution assay

A single colony was inoculated with the transformed cultures of TDP-43 and FUS in 500µl of media of SD-Ura and allowed to grow overnight on a shaking incubator at 30°C. Raf-Ura (YNB, raffinose and amino acid mixture lacking Tryptophan and Uracil) washes were given, i.e., dextrose in the medium was replaced by raffinose, and their O.D_600_. was adjusted to 0.2 and allowed to grow again in the same conditions. Following day, the O.D was adjusted to 0.2, and the culture was diluted up to 1/10000^th^ dilution (10^0^ to 10^-5^dilutions) and 3-6 µl of each dilution was spotted on to SD-Ura and SGal-Ura (galactose, YNB, Agar and amino acid mixture lacking Tryptophan and Uracil) plates and incubated at 30^0^C for 2-4 days, as all the plasmids used are galactose inducible.

For the double transformation of ALS mutant strains, two clones were selected and inoculated into an SD-Trp-Ura medium. Further plated on SD-Trp-Ura and SGal-Trp-Ura plates with control luciferase.

#### 4.2.4 Growth Assay

Transformed colonies of double transformants were inoculated in SD-Trp-Ura (Sucrose, YNB, and amino acid lacking mixture Uracil) medium overnight on a shaking incubator at 30°C. The next day, Raf-Trp-Ura (Raffinose, YNB, and amino acid lacking mixture Tryptophan Uracil) washes were given, and their O.D_600_. was adjusted to 0.2 and 20% galactose was added to each culture of Raf-Trp-Ura and these were added to 96 well plates in triplicates with blank (Raf-Trp-Ura + Gal). 96-well plate was placed in an Epoch2 BioTek Spectrophotometer for 48 hours at 30°C with continuous orbital shaking, and O.D_600_. was measured every 30 min. The data obtained after 48 hrs were plotted in GraphPad software, and the Growth curve was plotted for each culture.

#### 4.2.5 Microscopy

By using confocal microscopy, we analysed the protein aggregation status of the yeast cells as the FUS and TDP43 constructs had a GFP tag.All transformed *S. cerevisiae* strains were grown overnight in SD-Trp-Ura broth at 30°C. The next day, they were subjected to Raf-Trp-Ura^-^wash for the removal of residual dextrose. O.D_600_ was adjusted to 0.2 in Raf-Trp-Ura^-^. After 2-3 hours, they were induced with 20% galactose. The next day, they were centrifuged and resuspended in the residual media, and a smear was prepared and observed under a confocal microscope under a 60x objective and GFP laser. Images were processed using Fiji software.

#### 4.2.5 Quantitation of Image Analysis

Mutant strains transformed with different chaperones displayed two different morphologies, punctae and diffuse cytoplasmic signal. These were counted in 3 biological replicates for each strain. Chaperones that show more of the diffuse cytoplasmic expression and fewer punctae compared to luciferase are classified as suppressors, and chaperones that show a greater number of punctae and less of the diffuse cytoplasmic signal are classified as enhancers.

## 5.0 Acknowledgment

We would like to thank Amitabha Majumdar for providing the plasmids and the instrument setup required for this study. We would like to thank Meghal Desai for the plasmid library construction.

This work was supported by research grants from the Department of Biotechnology (DBT-IYBA) (BT/09/IYBA/2015/03) and the DBT RLS (BT/RLF/Re-entry/54/2013) to TB. SA recipients of the Indian Council of Medical Research (ICMR) senior research fellowship (SRF) (5/3/8/89/ITR-F/2020-ITR).

## 6.0 Authors contribution

The experiments are designed in this study by TB. The manuscript was written by SA and TB. Experiments were done by SA.

## 8.0 Declaration of generative AI in scientific writing

No generative AI is used during scientific writing

## Abbreviations

ALS: Amyotrophic Lateral Sclerosis
TDP-43: TAR DNA binding protein 43
FUS: Fused in Sarcoma
HSPs: Heat shock proteins
GFP: Green fluorescent protein

## 12.0 Supplementary figures

**Supplementary Fig S1. TDP-43 and FUS plasmid.**

ALS plasmid TDP-43 and FUS sequence map.

**Supplementary Fig S2. ALS mutants TDP-43 and FUS and their effect on toxicity in yeast cells.**

**S2a.** Spot dilution of plasmids TDP-43 and FUS with control after transformation in yeast.

**S2b.** 48hrs growth curve of TDP-43 and FUS, obtaining O.D. after 30 min at 600 nm (n=3, P<0.05, paired t-test).

**S2b.** Confocal Microscopy of GFP-tagged plasmids TDP-43 and FUS with control.

**Supplementary Fig S3. DnaJ domain chaperones overexpression library construction.**

DnaJ domain chaperones overexpression library was created from flies, which are homologous to the DnaJ domain proteins of humans, by Gateway cloning.

**Supplementary Fig S4. Effect of mutant TDP-43 and DnaJ chaperones on growth**

**S4a.** Enhancers and suppressors of TDP-43 are characterized based on the spot dilution assay on the control plate SD-Trp-Ura.

**S4c.** Weak enhancer and suppressor classification based on spot dilution assay of TDP-43 chaperones on control SD-Trp-Ura.

**Supplementary Fig S5. Effect of mutant FUS and DnaJ chaperones on growth**

**S5a.**Enhancers and suppressors of FUS are characterized based on the spot dilution assay on the control plate SD-Trp-Ura.

**S4c.**Weak enhancer and suppressor classification of FUS chaperones on control SD-Trp-Ura.

**Supplementary Fig S6. Phylogenetic tree for Hsp40 chaperones**

A phylogenetic tree was created using the Mega12 tool by using multiple sequence alignment of chaperones, showing region of protein conserved across species.

**Supplementary Table 1 Plasmid genotype and yeast strains.**

In supplementary Table1 genotype of yeast strain, TDP-43 and FUS plasmid as well as chaperone genotype, were mentioned.

**Supplementary Table 2 DnaJ domain chaperones and their related functions**

In this table, mentioned about homologue/orthologue of the *Drosophila* DnaJ domain chaperone as well as their predicted functions.

