## Supplementary material for "Overexpression of DnaJ chaperones in ameliorating toxicities associated with FUS and TDP-43": Supplemetary Fig

### Supplementary Fig S1

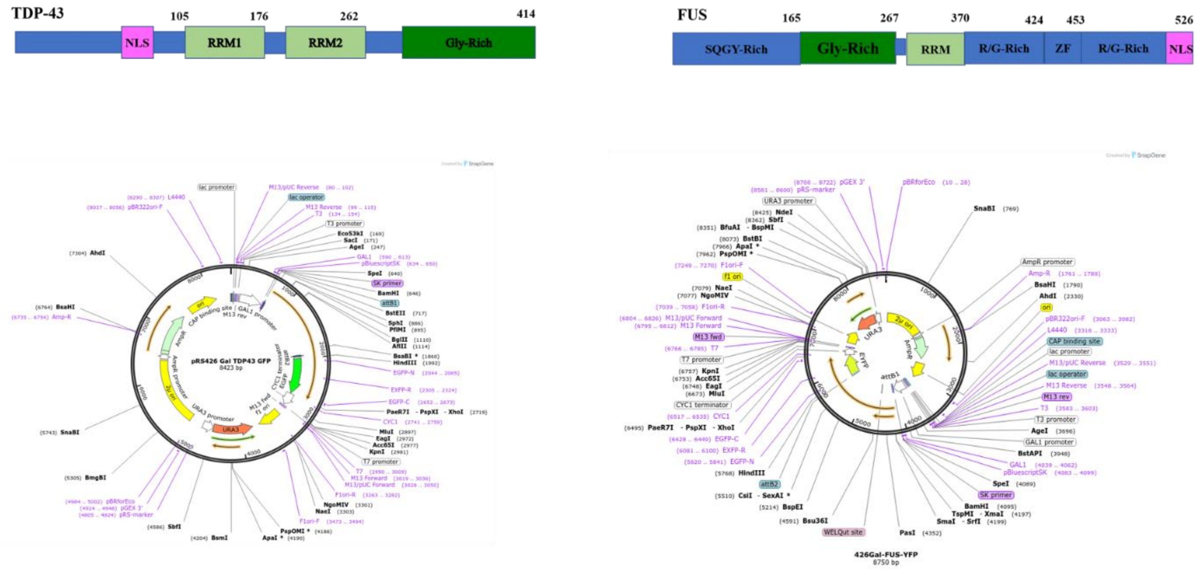


### Supplementary Fig S2

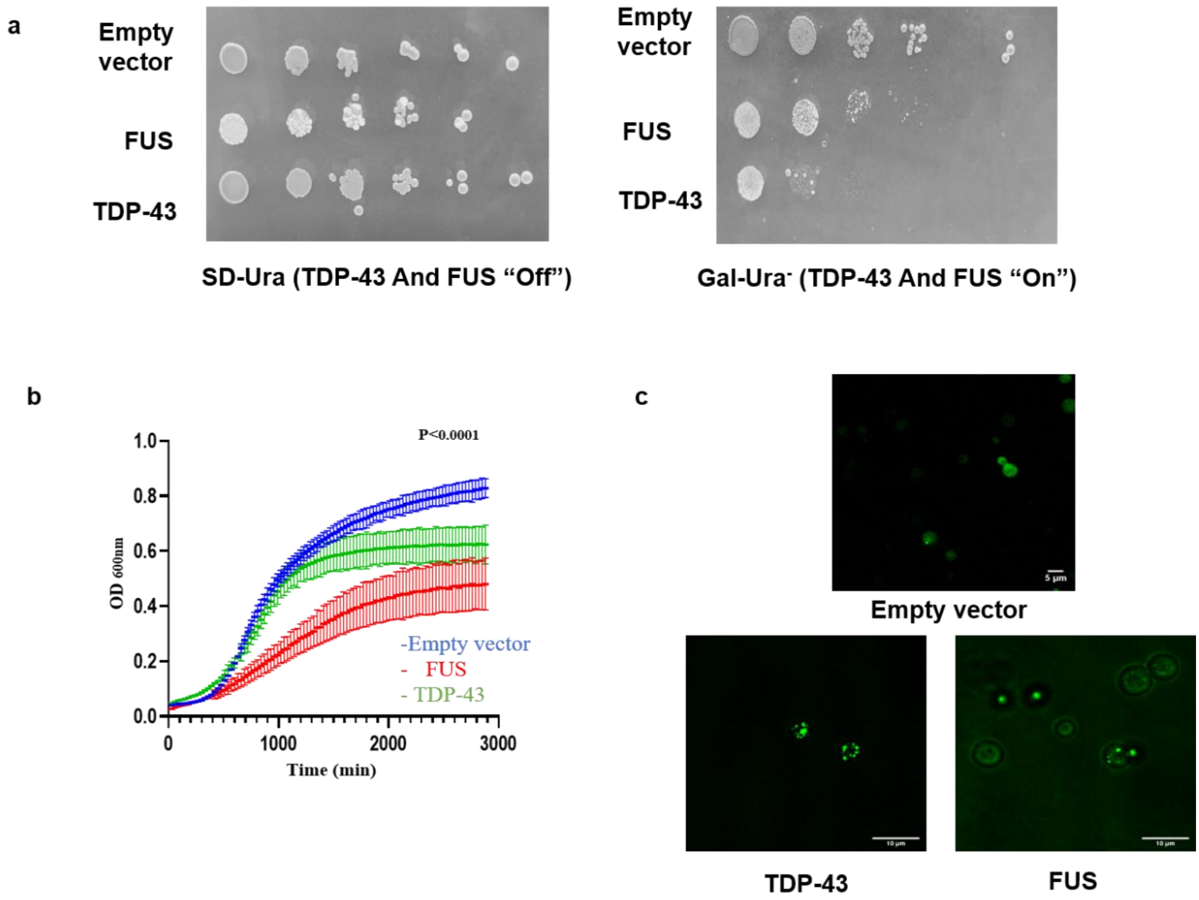

### Supplementary Fig S3

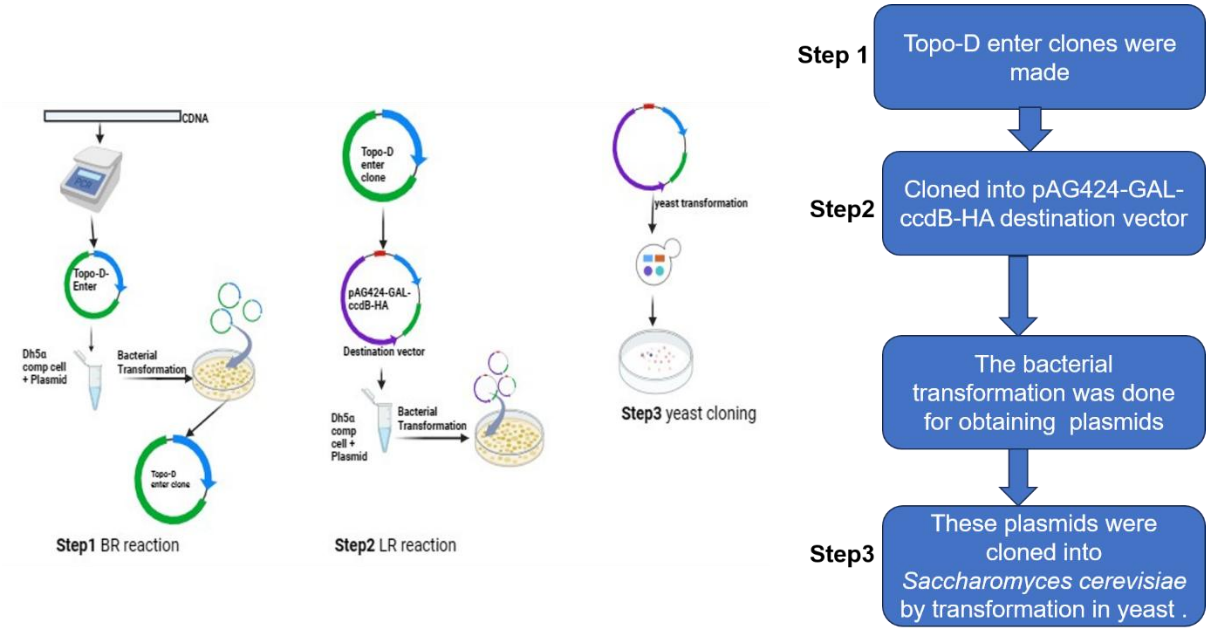


### Supplementary Fig S4

**a**

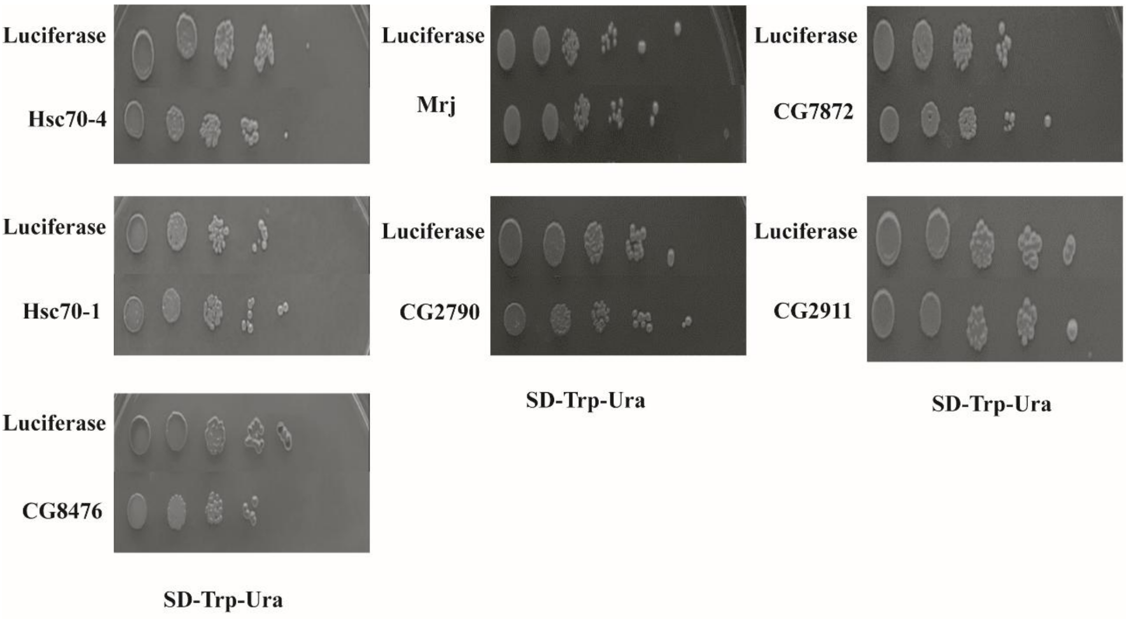

c

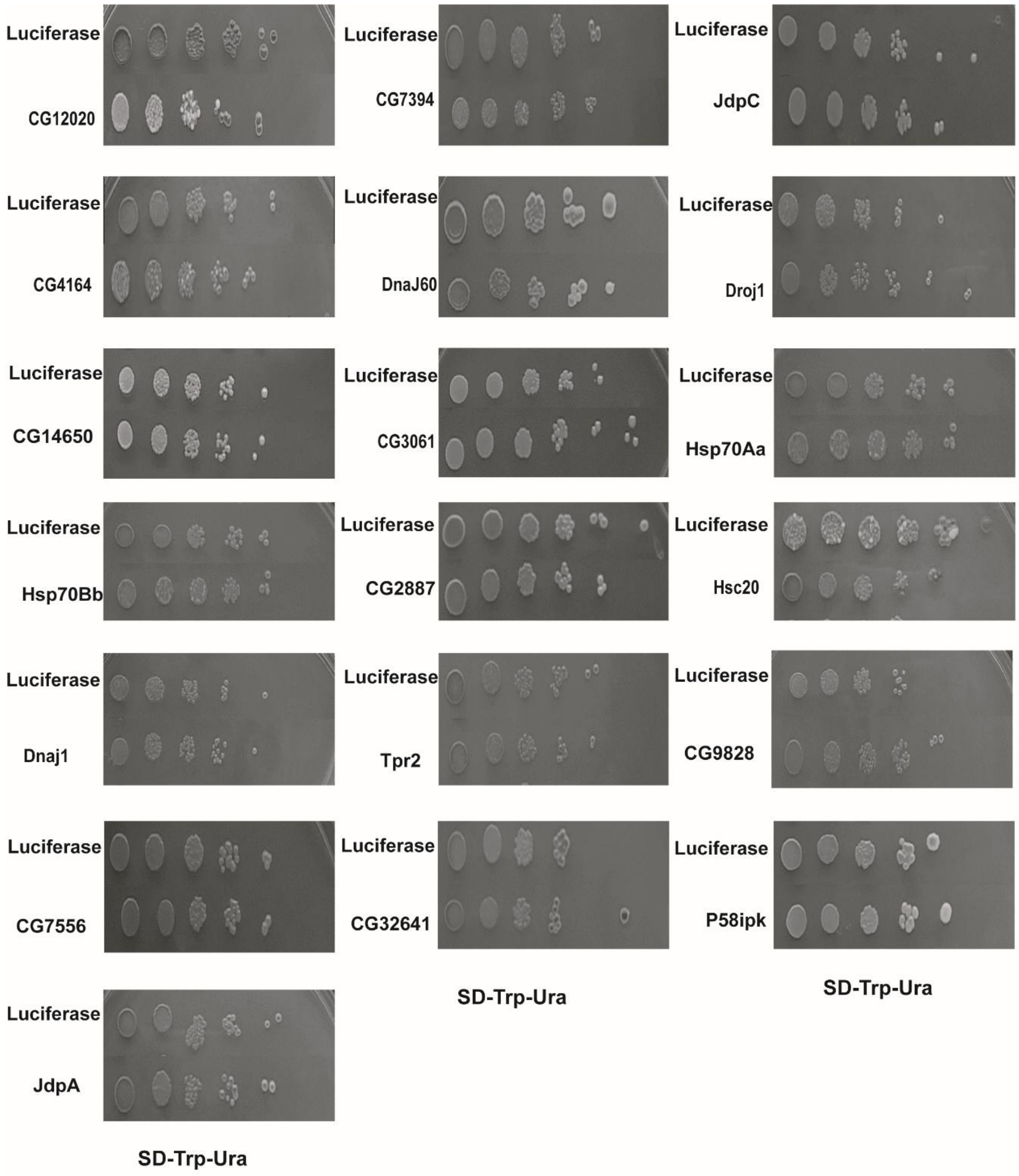
Luciferase
CG12020
Luciferase
CG7394
Luciferase
JdpC
Luciferase
CG4164
Luciferase
DnaJ60
Luciferase
Droj1
Luciferase
CG14650
Luciferase
CG3061
Luciferase
Hsp70Aa
Luciferase
Hsp70Bb
Luciferase
CG2887
Luciferase
Hsc20
Luciferase
Dnaj1
Luciferase
Tpr2
Luciferase
CG9828
Luciferase
CG7556
Luciferase
CG32641
Luciferase
P58ipk
Luciferase
JdpA
SD-Trp-Ura
SD-Trp-Ura
SD-Trp-Ura

### Supplementary Fig S5

**a**

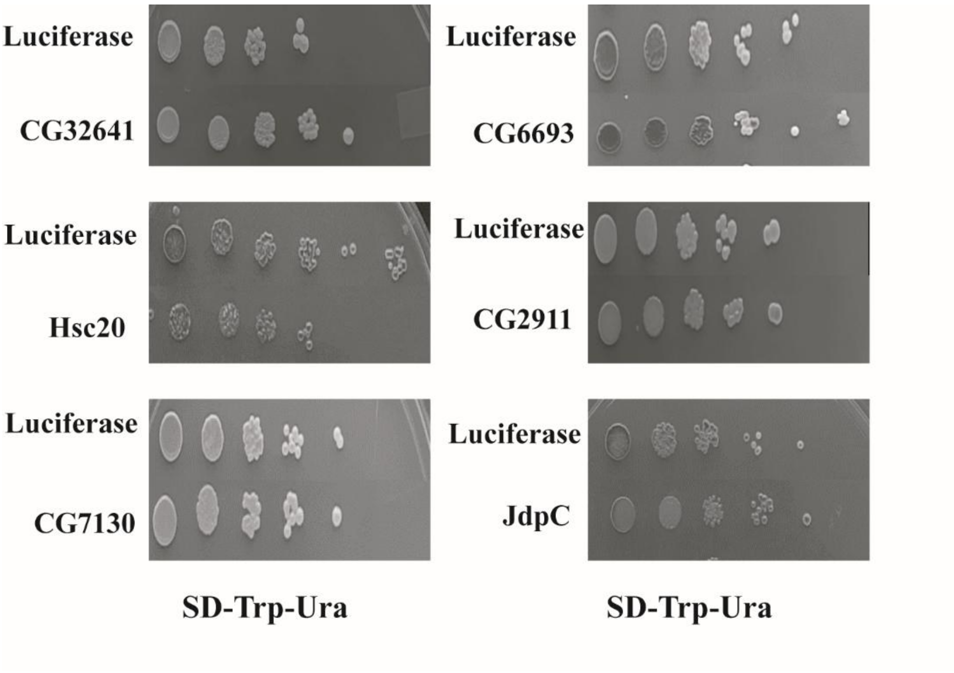

c

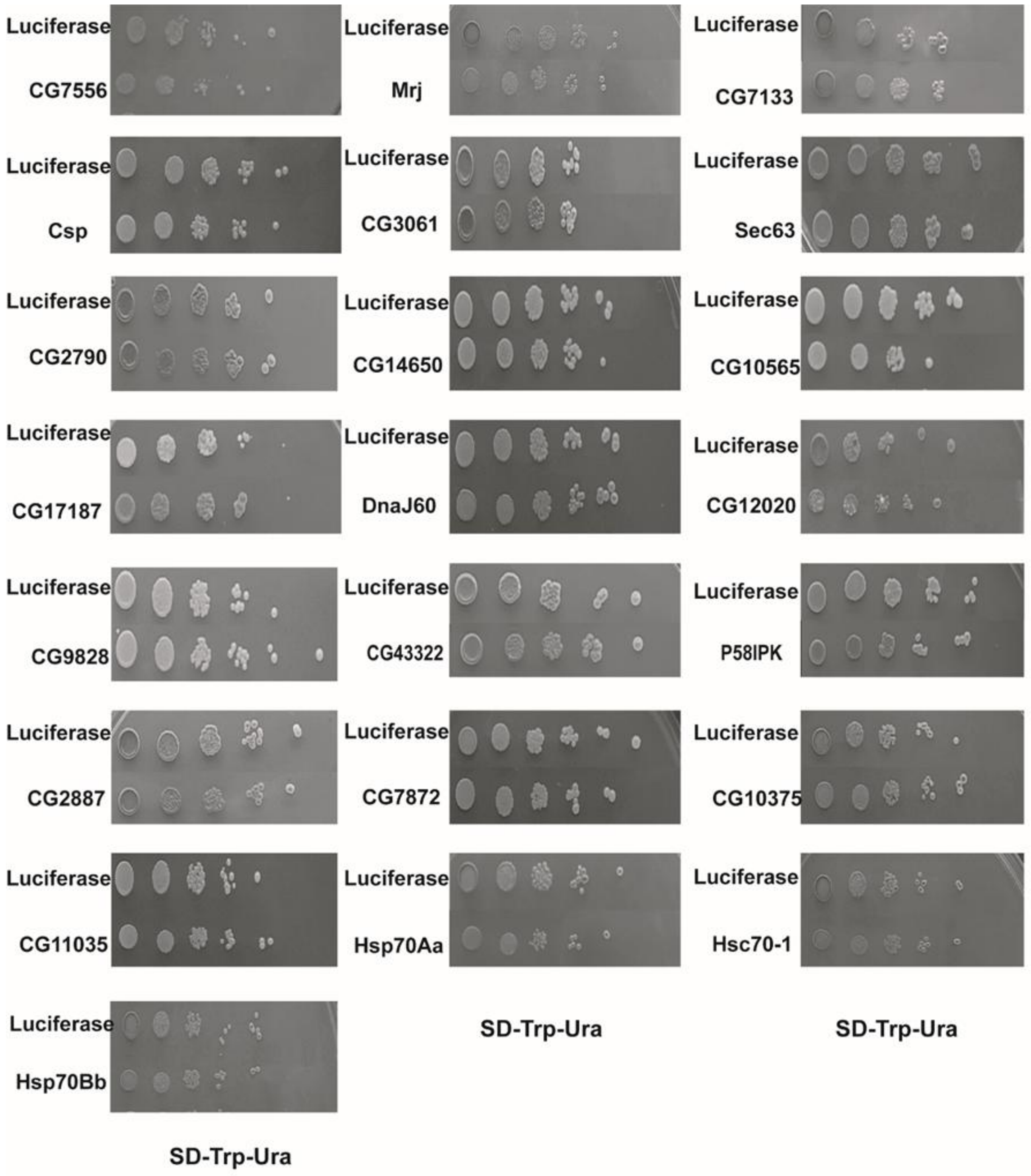
Luciferase
CG7556
Luciferase
Mrj
Luciferase
CG7133
Luciferase
Csp
Luciferase
CG3061
Luciferase
Sec63
Luciferase
CG2790
Luciferase
CG14650
Luciferase
CG10565
Luciferase
CG17187
Luciferase
DnaJ60
Luciferase
CG12020
Luciferase
CG9828
Luciferase
CG43322
Luciferase
P58IPK
Luciferase
CG2887
Luciferase
CG7872
Luciferase
CG10375
Luciferase
CG11035
Luciferase
Hsp70Aa
Luciferase
Hsc70-1
Luciferase
Hsp70Bb
SD-Trp-Ura
SD-Trp-Ura
SD-Trp-Ura

**Supplementary Fig S6**

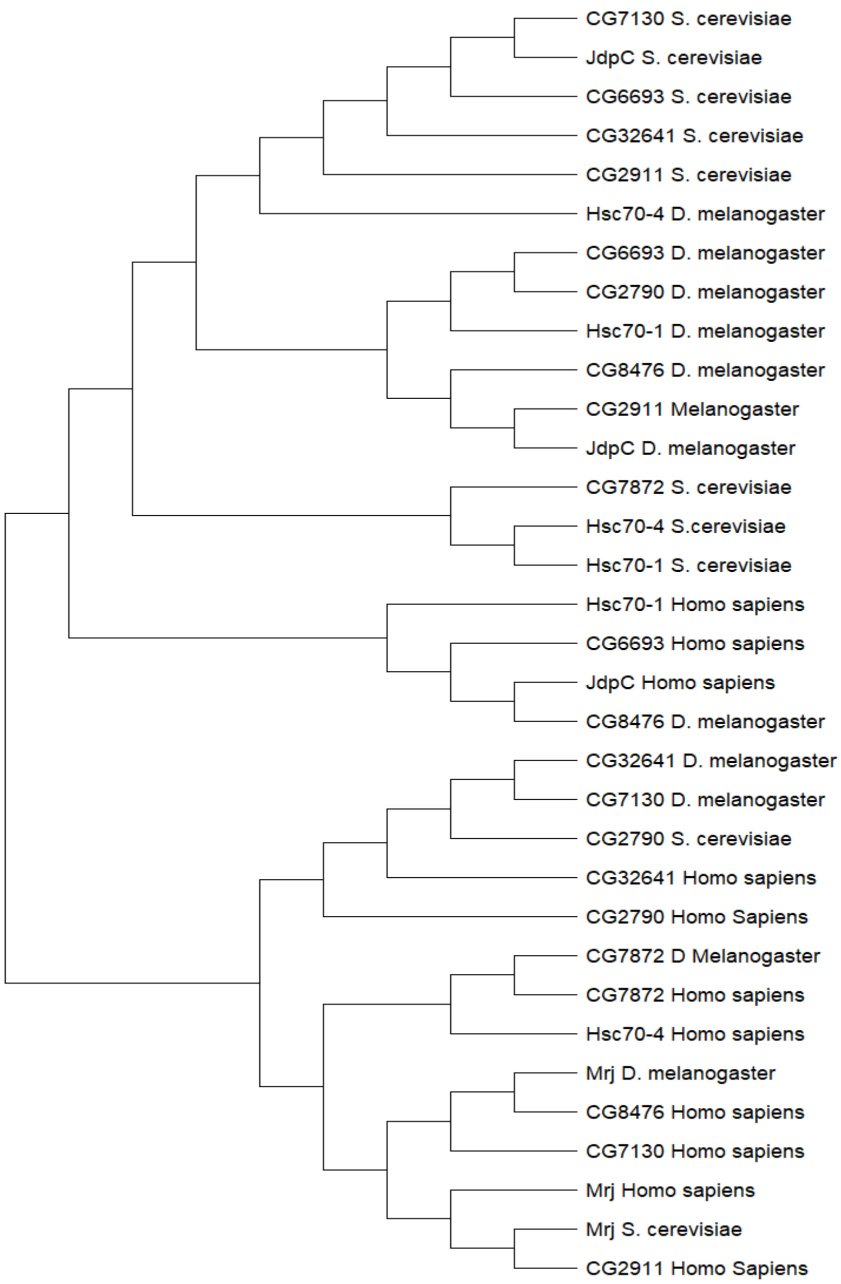

**Supplementary Table S1**

| Sr. No. | Yeast Strains Used | Genotype |
|---|---|---|
| 1 | BY4741 | MATa his3Δ1 leu2Δ0 met15Δ0 ura3Δ0 |
| 2 | BY4742 | MATα his3Δ1 leu2Δ0 lys2Δ0 ura3Δ0 |
| 3 | W303 | MATa/MATα {leu2-3,112 trp1-1 can1-100 ura3-1 ade2-1 his3-11,15} [$phi^+$] |
| | **Plasmids** | |
| 1 | CG27467 | pRS426 Gal TDP43 GFP p |
| 2 | CG29592 | 426 Gal-FUS-YFP |
| | **DnaJ Chaperones** | |
| 1 | CG7872 | pAG424Gal-CG7872/Trp |
| 2 | Mrj | pAG424Gal-Mrj/Trp |
| 3 | Hsc70-4 | pAG424Gal-Hsc70-4/Trp |
| 4 | Hsc70-1 | pAG424Gal-Hsc70-1/Trp |
| 5 | CG2790 | pAG424Gal-CG2790/Trp |
| 6 | CG32641 | pAG424Gal-CG32641/Trp |
| 7 | CG6693 | pAG424Gal-CG6693/Trp |
| 8 | JdpC | pAG424Gal-JdpC/Trp |
| 9 | CG8476 | pAG424Gal-CG8476/Trp |
| 10 | JdpA | pAG424Gal-JdpA/Trp |
| 11 | CG4164 | pAG424Gal-CG4164/Trp |
| 12 | Hsc20 | pAG424Gal-Hsp40/Hsp70/Trp |
| 13 | CG2911 | pAG424Gal-CG2911/Trp |
| 14 | CG3061 | pAG424Gal-CG3061/Trp |
| 15 | Tpr2 | pAG424Gal-Tpr2/Trp |
| 16 | CG14650 | pAG424Gal-CG14650/Trp |
| 17 | Hsp70Bb | pAG424Gal-Hsp70Bb/Trp |
| 18 | Hsp70Aa | pAG424Gal-Hsp70Aa/Trp |
| 19 | CG12020 | pAG424Gal-CG12020/Trp |
| 20 | CG7394 | pAG424Gal-CG7394/Trp |
| 21 | DnaJ60 | pAG424Gal-DnaJ60/Trp |
| 22 | DnaJ1 | pAG424Gal-DnaJ1/Trp |
| 23 | Droj1 | pAG424Gal-Droj1/Trp |
| 24 | CG9828 | pAG424Gal-CG9828/Trp |
| 25 | P58IPK | pAG424Gal-P58IPK/Trp |
| 26 | CG2887 | pAG424Gal-CG2887/Trp |
| 27 | CG7130 | pAG424Gal-CG7130/Trp |
| 28 | Csp | pAG424Gal-Csp/Trp |
| 29 | Sec63 | pAG424Gal-Sec63/Trp |
| 30 | CG10565 | pAG424Gal-CG10565/Trp |
| 31 | CG17187 | pAG424Gal-CG17187/Trp |

| | | |
|---|---|---|
| 32 | CG43322 | pAG424Gal-CG43322/Trp |
| 33 | CG10375 | pAG424Gal-CG10375/Trp |
| 34 | CG11035 | pAG424Gal-CG11035/Trp |
| 35 | CG7133 | pAG424Gal-CG7133/Trp |
| 36 | CG30156 | pAG424Gal-CG30156/Trp |
| 37 | CG5001 | pAG424Gal-CG5001/Trp |
| 38 | CG7556 | pAG424Gal-CG7556/Trp |

**Supplementary Table S2**

| Sr.no | Gene name | Homologue/Orthologue | Predicted Function |
|---|---|---|---|
| 1 | CG4164 | DNAJB11 | Encodes a secreted protein that activates integrin signaling. It is involved in synapse maturation regulation and stem cell maintenance |
| 2 | CG2887 | DNAJB5 | Protein-folding chaperone binding activity and unfolded protein binding activity. Predicted to be involved in chaperone cofactor-dependent protein refolding |
| 3 | CG8476 | DNAJB9 | Enable misfolded protein binding activity and protein-folding chaperone binding activity. Predicted to be involved in ubiquitin-dependent ERAD pathway. Predicted to be active in endoplasmic reticulum |
| 4 | CG6693 | DNAJC9 | Enable heat shock protein binding activity. Predicted to be active in cytoplasm and nucleus. Is expressed in several structures, including CNS glial cell; ectoderm; embryonic central brain neurons; embryonic/larval nervous system; and extended germ band embryo |
| 5 | CG11035 | DNAJC30 | Predicted to be involved in brain development and regulation of mitochondrial ATP synthesis coupled proton transport. Predicted to be located in mitochondrion. Is expressed in adult head. |
| 6 | CG7872 | DNAJC25 | Involved in protein folding. |
| 7 | Mrj | DNAJB6 | Enable protein folding chaperones activity and unfolded protein activity. |
| 8 | CG12020 | DNAJB13 | Protein folding chaperones activity, and unfolded protein activity, chaperone cofactor-dependent protein refolding. |
| 9 | DnaJ1 | DNAJB13 | Enable protein-folding chaperone binding activity and unfolded protein binding activity. Predicted to be involved in chaperone cofactor-dependent protein refolding |
| 10 | DnaJ60 | DNAJC4 | Unfolded protein binding. It is involved in the biological process described with: spermatogenesis; protein refolding |
| 11 | CG2911 | DNAJC24 | Enable ATPase activator activity |

| | | | |
|---|---|---|---|
| 12 | CG5001 | DNAJB5 | Enable protein-folding chaperone binding activity and unfolded protein binding activity. Predicted to be involved in chaperone cofactor-dependent protein refolding and predicted to be active in cytosol |
| 13 | CG9828 | DNAJA2 | Enable ATP binding, Enable Hsp70 protein binding, protein binding, and protein folding chaperones. |
| 14 | CG3061 | DNAJB12 | Involved in cellular response to misfolded protein and chaperone cofactor-dependent protein refolding. |
| 15 | CG32641 | DNAJB9 | It is involved in the biological process described with: cellular response to misfolded protein; chaperone cofactor-dependent protein refolding; ubiquitin-dependent ERAD pathway. |
| 16 | CG17187 | DNAJC17 | Predicted to enable nucleic acid binding activity. Predicted to be involved in spliceosome complex disassembly. Part of precatalytic spliceosome. |
| 17 | CG7394 | DNAJC19 | Predicted to enable ATPase activator activity. Predicted to be involved in protein import into mitochondrial matrix. Predicted to be part of PAM complex, Tim23 associated import motor |
| 18 | Hsc20 | HSCB | A mitochondrial chaperone that facilitates iron-sulfur cluster assembly or transfer from the product of IscU to other scaffold proteins. Hsc20 is required for larval growth and cellular iron sensing. |
| 19 | P58IPK | DNAJC3 | Predicted to enable misfolded protein binding activity and protein-folding chaperone binding activity. Predicted to be involved in protein folding in endoplasmic reticulum. Located in endomembrane system. Is expressed in several structures, including embryonic/larval digestive system |
| 20 | CG7556 | DNAJC1 | Located in endomembrane system. Is expressed in adult head; embryonic/larval salivary gland; embryonic/larval salivary gland body |
| 21 | CG14650 | DNAJC14 | Its molecular function is unknown. The biological processes in which it is involved are not known |
| 22 | Droj1 | Hsap\DNAJB5 | Its molecular function is described by: protein binding; unfolded protein binding; protein-folding chaperone binding; DNA-binding transcription factor binding. |
| 23 | Tpr2 | DNAJC7 | Molecular function is described by: heat shock protein binding |

| | | | |
|---|---|---|---|
| 24 | Hsc70-1 | HSPA2 | Enable ATP hydrolysis activity; heat shock protein binding activity; and protein folding chaperone. Predicted to be involved in chaperone cofactor-dependent protein refolding and protein refolding. Predicted to be active in cytosol; nucleus; and plasma membrane. |
| 25 | Hsc70-4 | HSPA8 | Enables protein-folding chaperone binding activity. Involved in several processes, including RNA-mediated gene silencing; axon development; and late endosomal micro autophagy. |
| 26 | Hsp70Aa | Hsap\HSPA1B | To enable ATP hydrolysis activity; heat shock protein binding activity; and protein folding chaperone. Involved in heat shock-mediated polytene chromosome puffing and response to hypoxia. |
| 27 | Hsp70Bb | Hsap\HSPA1A | Encodes a protein involved in response to hypoxia heat shock |
| 28 | CG2790 | DNAJC21 | Predicted to allow nucleic acid binding and zinc ion binding activity. Active in cytoplasm |
| 29 | JdpC | DNAJC12 | Allow protein folding chaperone binding activity and unfolded protein binding |
| 30 | JdpA | - | - |
| 31 | Csp | DNAJC5 | Cysteine string protein (CSP) encodes a synaptic vesicle-associated co-chaperone of Hsc70 that is vital for presynaptic proteostasis and maintenance of synaptic function. |
| 32 | CG30156 | DNAJB12 | Involved in cellular response to misfolded protein and chaperone cofactor-dependent protein refolding. Predicted to be active in endoplasmic reticulum membrane |
| 33 | CG7387 | DNAJA3 | Predicted to enable heat shock protein binding activity and unfolded protein binding activity. Predicted to be involved in mitochondrion organization and negative regulation of apoptotic process |
| 34 | CG8531 | DNAJC11 | Enable ATPase activator activity. Predicted to be involved in cristae formation and protein import into the mitochondrial matrix |
| 35 | CG10565 | DNAJC2 | Enable Hsp70 protein binding activity and ribosome binding activity. Predicted to be involved in 'de Novo cotranslational protein folding. |
| 36 | Sec63 | DNAJA3 | Predicted to enable protein transmembrane transporter activity. Predicted to be involved in SRP-dependent translational protein targeting to membrane and post-translational protein targeting to endoplasmic reticulum membrane. Located in fusome. |
| 37 | CG7130 | - | - |

| | | | |
|---|---|---|---|
| 38 | CG43322 | DNAJC28 | Predicted to be involved in Golgi organization; Golgi vesicle prefusion complex stabilization; and retrograde vesicle-mediated transport, Golgi to endoplasmic reticulum |
